# CAS-LiveFISH: Simple and versatile imaging of genomic loci in live mammalian cells and early pre-implantation embryos

**DOI:** 10.1101/2020.08.25.265306

**Authors:** Yongtao Geng, Alexandros Pertsinidis

**Affiliations:** Structural Biology Program, Memorial Sloan Kettering Cancer Center, New York, NY 10065, USA

## Abstract

Visualizing the 4D genome in live cells is essential for understanding its regulation. Programmable DNA-binding probes, such as fluorescent clustered regularly interspaced short palindromic repeats (CRISPR) and transcription activator-like effector (TALE) proteins have recently emerged as powerful tools for imaging specific genomic loci in live cells. However, many such systems rely on genetically-encoded components, often requiring multiple constructs that each must be separately optimized, thus limiting their use. Here we develop efficient and versatile systems, based on *in vitro* transcribed single-guide-RNAs (sgRNAs) and fluorescently-tagged recombinant, catalytically-inactivated Cas9 (dCas9) proteins. Controlled cell delivery of pre-assembled dCas9-sgRNA ribonucleoprotein (RNP) complexes enables robust genomic imaging in live cells and in early mouse embryos. We further demonstrate multiplex tagging of up to 3 genes, tracking detailed movements of chromatin segments and imaging spatial relationships between a distal enhancer and a target gene, with nanometer resolution in live cells. This simple and effective approach should facilitate visualizing chromatin dynamics and nuclear architecture in various living systems.

## INTRODUCTION

The dynamic genome organization in the nucleus directs many important processes, such as gene expression regulation, epigenetic inheritance, recombination, DNA replication and genome maintenance, in normal physiology and in disease. Fluorescence *in-situ* hybridization (FISH) methods have been widely used to detect specific DNA sequences, providing a wealth of information on 3D genome architecture. However FISH relies on chemical fixation, precluding tracking real-time dynamics in live cells. Moreover, harsh treatments with high heat and denaturants might compromise the integrity of genomic structures, especially on the kilobase and nanometer scales. To address some of these limitations, fluorescent programmable DNA binding proteins have emerged as a promising tool for visualizing specific genomic loci in live cells. Fluorescent TALE proteins, engineered to recognize specific DNA sequences, such as major satellite and telomere repeats, were used to visualize repetitive DNA sequences in live human and mouse cells as well as in fly and mouse embryos (Ma et al., 2013; Miyanari et al., 2013; Yuan et al., 2014). Although engineering a separate TALE protein is required for each genomic locus, the use of CRISPR-Cas9 proteins, whose DNA recognition can be programmed by a single-guide RNA, significantly simplified targeting (Anton et al., 2014; Chen et al., 2013). Multi-color approaches based on orthogonal dCas9 proteins (Chen et al., 2016; Ma et al., 2015), or by fusing various RNA protein binding motifs to the gRNA (Fu et al., 2016; Ma et al., 2016; Qin et al., 2017), have also been demonstrated.

Most of the previous CRISPR imaging applications focused on visualizing repetitive DNA elements, where the signal amplification from multiple bound dCas9 molecules facilitates detection. Although significantly more challenging, dCas9 can also be used to tag non-repetitive DNA elements, when programmed with multiple gRNAs that target a specific locus in tandem (Chen et al., 2013; Gu et al., 2018; Maass et al., 2018). However, robust tagging for non-repetitive locus imaging can be challenged by low efficiencies. Often the system needs to be engineered to optimize expression levels of fluorescent dCas9 as well as gRNAs. This issue is further exacerbated by the additional complexity of multi-color systems, which use multiple expression constructs, each one needing separate optimization, limiting the broad use of CRISPR for genome imaging applications.

Here we report a simple and versatile CRISPR imaging system based on *in vitro* transcribed gRNAs and fluorescent recombinant dCas9 proteins. Controlled cell delivery enables straightforward optimization of the system to achieve robust tagging and high signal-to-noise (SNR) imaging. Simultaneous delivery of multiple gRNAs is also simplified, allowing tandem tagging of endogenous loci. Using this approach, CRISPR/Cas9-mediated *in situ* tagging of genomic loci in live cells, CAS-LiveFISH, we achieve simultaneous multi-color tagging of up to 3 genes and efficient tagging of low-repeat-containing and non-repetitive loci. We further show that pre-assembled RNPs can successfully visualize genomic loci in live pre-implantation mouse embryos, as a proof-of-principle for CAS-LiveFISH imaging in more complex living systems. Finally, we demonstrate tracking of sub-diffusive motions of chromatin loci and 2-color nanometer distance measurements between a distal enhancer and a target gene in live cells. These results illustrate the efficiency, versatility and capability of CAS-LiveFISH for imaging genome organization and dynamics.

## RESULTS

### Imaging of genomic loci by delivery of *in vitro* transcribed gRNAs and recombinant fluorescent dCas9

We first test the ability of *in vitro* transcribed gRNAs to target cell-expressed dCas9-EGFP to specific genomic loci (Figure 1**A**). We deliver gRNAs targeting telomeres in live U-2 OS cells using electroporation, together with a plasmid encoding for BFP-LifeAct to assist identification of transfected cells. Telomeres are visible in transfected cells, while a control gRNA (sgGal4) does not result in any nuclear puncta, demonstrating the specificity of the approach (Figure 1**B**, Supplementary Figure 1**A**). As an alternative, *in vitro* transcribed gRNAs are microinjected in live U-2 OS cells, together with Alexa 647-Benzylguanine, as a nuclear “counter-stain” for SNAP-tagged RNA Pol II. Like electroporation, microinjection of gRNAs results in robust dCas9-EGFP nuclear puncta (Figure 1**C**), suggesting efficient tagging of telomeres and illustrating versatility among cell delivery approaches.

**Figure 1.**
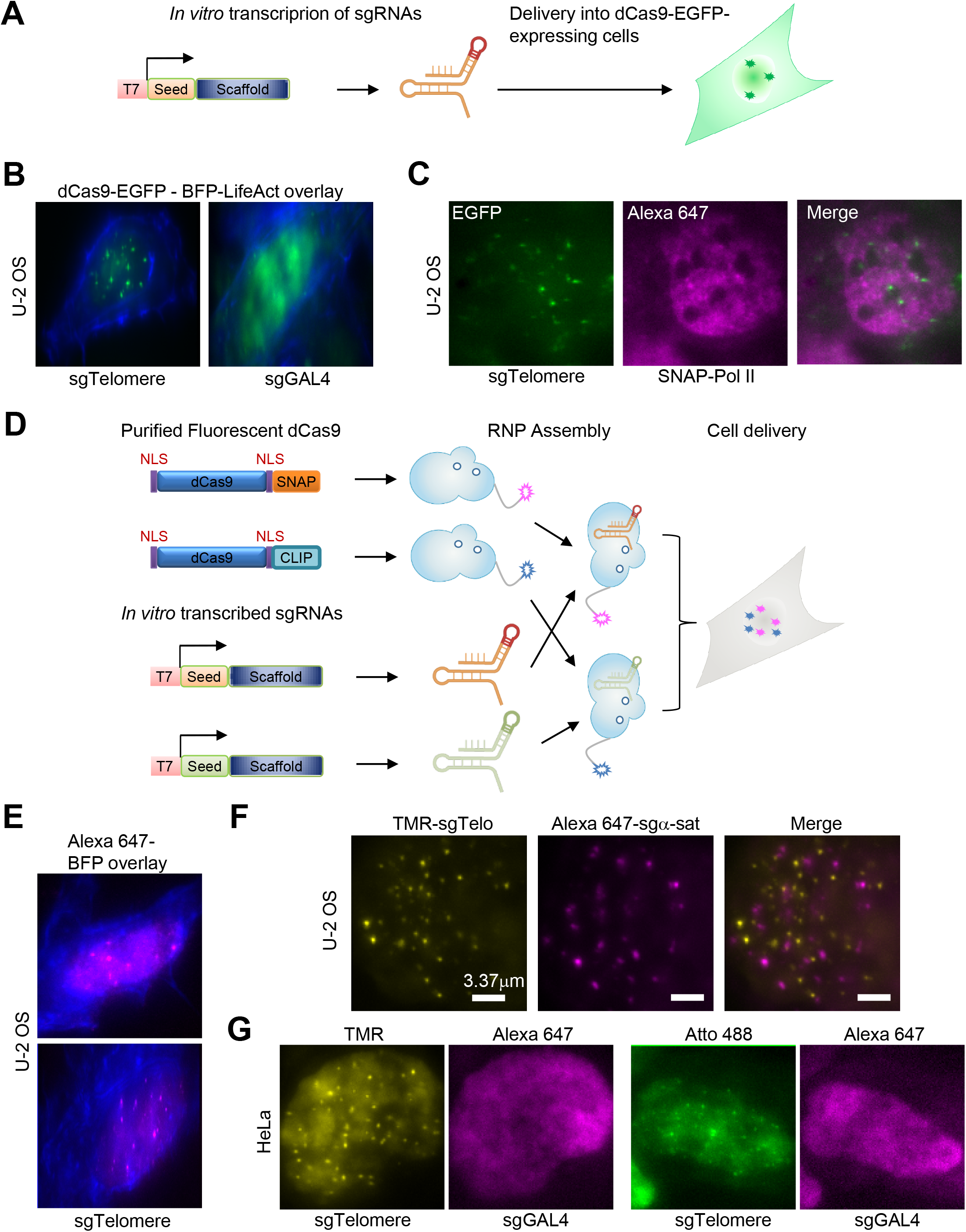
Imaging genomic loci in live cells with CAS-LiveFISH. (**A**) Schematic of cell delivery of *in vitro* transcribed gRNAs. (**B, C**) Visualization of telomeres in U-2 OS cells stably expressing dCas9-EGFP, by electroporation (**B**) or microinjection (**C**) of an *in vitro* transcribed gRNA (sgTelomere). A non-targeting control (sgGal4) shows no discernible nuclear puncta (**B**), demonstrating locus-specific tagging. In (**B**), cells are imaged 20hrs after electroporation. Co-injection of a BFP-LifeAct-expressing plasmid (**B**) or Alexa 647-Benzylguanine (**C**) results in simultaneous visualization of actin filaments and RNA Polymerase II (SNAP-Pol II), respectively. (**D**) Schematic of recombinant fluorescent dCas9:gRNA RNP assembly and cell delivery. (**E, F, G**) Visualization of genomic loci in U-2 OS (**E**) and HeLa (**F, G**) cells using pre-assembled dCas9-gRNA RNPs. (**F**) Co-delivery of TMR-RNPs targeting telomeres and Alexa 647-RNPs targeting a-satellite sequences shows distinct, non-overlapping nuclear puncta. (**G**) Co-injection of telomere-targeting Atto 488- or TMR-RNPs and non-targeting control Alexa 647-RNPs (sgGal4) shows no discernible nuclear puncta in the Alexa 647 channel, demonstrating locus-specific tagging and minimal cross-talk between colors.

Based on the results with *in vitro* transcribed gRNAs, we next test the ability of pre-assembled fluorescent dCas9-gRNA ribonucleoproteins (RNPs) to tag specific genomic loci (Figure 1**D**). We first prepare fluorescent dCas9-SNAPf and dCas9-CLIPf proteins (Supplementary Figure 2) and validate their *in vitro* DNA binding activity and specificity when assembled with *in vitro* transcribed gRNAs into RNPs (Supplementary Figure 3). We next test the preassembled RNPs in live cells. Delivery of Alexa 647-labelled RNPs targeting telomeres into U-2 OS cells by microinjection results in robust tagging, similar to that achieved by cell expressed dCas9-EGFP (Figure 1**E**). Co-delivery of two types of RNPs into live U-2 OS cells, TMR-labelled RNPs targeting telomeres and Alexa 647-labeled RNPs targeting α-satellite repeats, results in distinct, non-overlapping nuclear puncta (Figure 1**F**). To further test the specificity of tagging, we assemble Atto 488- or TMR-labelled RNPs targeting telomeres, and non-targeting (sgGal4) Alexa 647-labelled RNPs as a control. Co-delivery into live HeLa cells results in robust telomere tagging, visible in the Atto 488 or TMR channel, while no nuclear puncta are observable in the Alexa 647 channel (Figure 1**G**). Together, these results are consistent with *in vitro* competition experiments that show minimum cross-talk (Supplementary Figure 4**A, B**) and illustrate specificity in two-color genome tagging as well as versatility among cell types.

### CAS-LiveFISH imaging of genomic loci in live pre-implantation embryos

Although numerous demonstrations of CRISPR-based genomic imaging have been reported in cultured cells, applications in living organisms have been thus far limited. We reason that the versatility of the pre-assembled fluorescent RNP system could also be used to tag genomic loci in live mammalian pre-implantation embryos. We thus microinject preassembled TMR-labelled dCas9 RNPs targeting telomeres, into 2-cell stage mouse embryos (Figure 2**A**). Robust nuclear TMR puncta can be immediately discerned (Figure 2**B**), while a co-injected Alexa 647-Gal4 RNPs show only diffuse nuclear signal (Supplementary Figure 5**A**). TMR-tagged telomeres can be also observed at the 4-cell stage, ~1 day after microinjection (Figure 2**C**), after the second cleavage division, indicating that fluorescent RNPs are stable enough *in vivo* to potentially track dynamics over developmentally relevant time-scales in the early mouse embryo. These results show successful proof-of-principle use of the CAS-LiveFISH system for imaging genomic loci in live mammalian embryos.

**Figure 2.**
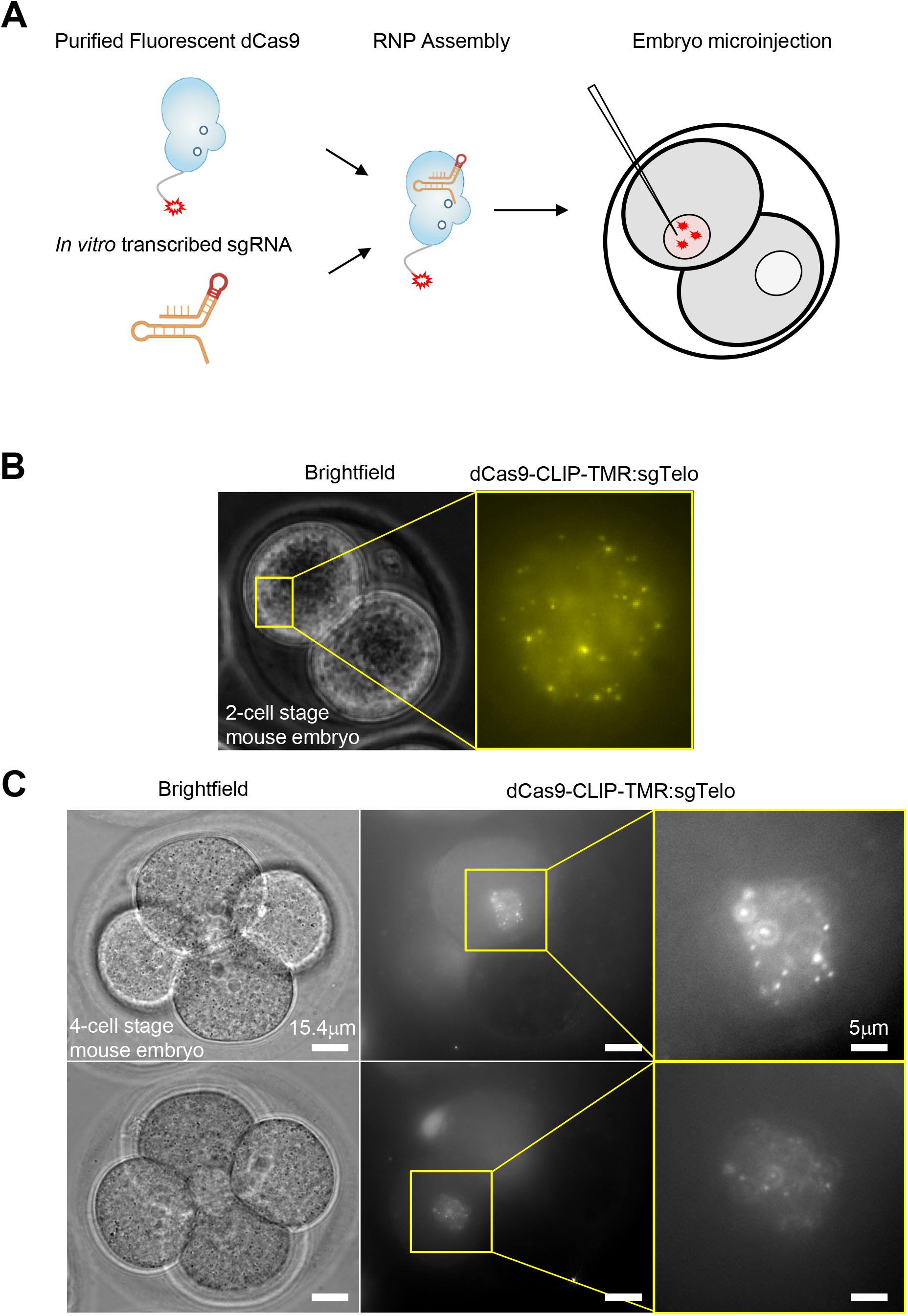
CAS-LiveFISH imaging in live mouse pre-implantation embryos. (**A**) Schematic of fluorescent RNP assembly and delivery into 2-cell stage embryos. (**B**) Brightfield and fluorescence images of a live 2-cell stage embryo, microinjected with TMR-sgTelo RNPs. Bright nuclear puncta are visible, indicating robust telomere labelling. (**C**) Brightfield and fluorescence images of a 4-cell stage embryo, one day after injecting one of the cells at the 2-cell stage with TMR-sgTelo RNPs. Nuclear puncta are visible in the two daughter nuclei. Top and bottom images correspond to two different focal planes, to bring each of the two nuclei in focus.

### Pools of pre-assembled RNPs enable visualizing the HPV-18 integration site in live HeLa cells

HeLa cells contain a genomic integration site of Human Papilloma Virus 18 (HPV-18) near the *MYC* oncogene (Figure 3**A**). HPV integration plays a prominent role in the initiation of cervical cancers (Bosch et al., 2002) and in HeLa cells the HPV integration site creates a powerful transcriptional enhancer that drives *MYC* activation (Adey et al., 2013). The HeLa HPV-18 integration site contains parts of the viral genome, as well as local integration-associated rearrangements of the host genome, with host and viral sequences amplified to copy numbers of ~4-30 (Adey et al., 2013).

**Figure 3.**
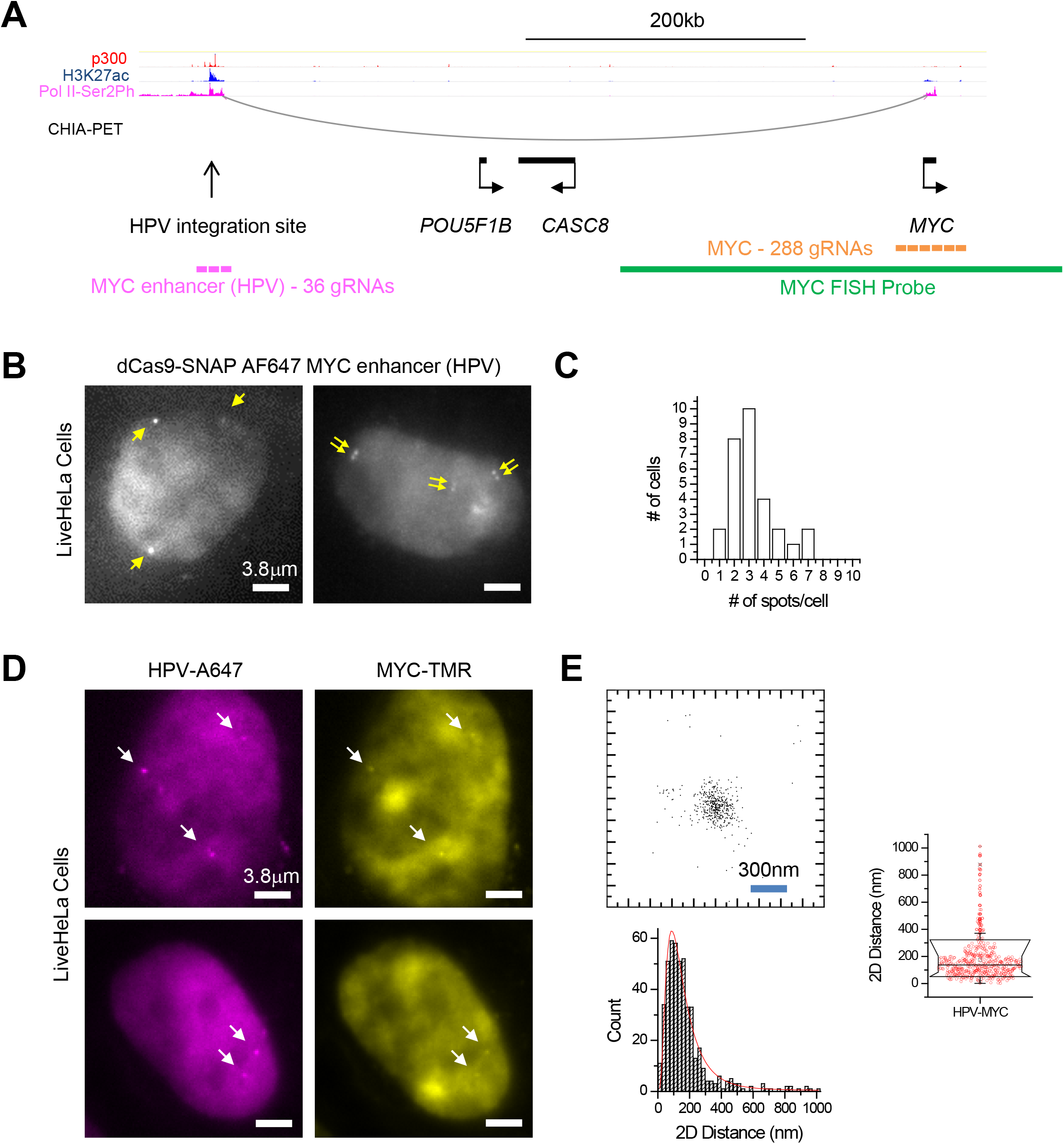
Pools of pre-assembled RNPs enable visualizing the *MYC* locus in live HeLa cells. (**A**) Organization of the extended *MYC* locus though proximity ligation (ChIA-PET) and ChIP-Seq assays. HeLa Pol II ChIA-PET data (GSM832461) (Li et al., 2012),as well as ChIP-Seq data for p300 (GSM935500), H3K27ac (GSM733684) and Pol2-Ser2Ph (GSM935383), are visualized using the WashU Epigenome Browser (Zhou et al., 2011). (**B**) Visualization of the HPV integration site using Alexa 647-labeled dCas9 RNPs, assembled with a pool of 36 distinct gRNAs. The right cell contains doublets of closely-spaced puncta, indicative of replicated loci in S/G2. (**C**) Distribution of number of discernible Alexa 647-HPV spots per cell. (**D**) Two-color imaging of HPV integration site (Alexa 647) and *MYC* (TMR) using co-delivery of preassembled RNPs in live HeLa cells. (**E**) 2D scatter plot of relative positions and statistics (histogram and box plot) of 2D distances between Alexa 647-HPV and TMR-MYC in live HeLa cells. 2D distance is 177±155 nm (mean±S.D.; *n*=472 measurements from >30 loci).

We design 36 distinct gRNAs targeting the HPV integration site and assemble them with Alexa 647 dCas9 into fluorescent RNPs. Delivery of this HPV-targeting pool of RNPs into live HeLa cells typically results in ~2-4 nuclear puncta per cell (Figure 3**B, C**). Occasionally we also resolve doublets in close proximity, suggestive of replicated loci in S/G2 (Chen et al., 2013). To verify the specificity of dCas9 tagging, we perform FISH with a probe labelling the extended *MYC* locus. Two-color imaging shows that the HPV-Alexa 647 puncta are within ~200-300 nm from the centroid of the *MYC* FISH puncta (Supplementary Figure 6**A, B**). These results demonstrate successful locus-specific tagging of the low-repeat containing HPV integration site, using pools of pre-assembled RNPs.

### Multiplex CAS-LiveFISH imaging of up to 3 genes

We reason that the minimal crosstalk between separately assembled two-color RNPs should also be extendable to higher multiplexing. We thus assemble Atto 488- and TMR-labeled dCas9 RNPs programmed with gRNAs targeting the *MUC1* and *MUC4* genes, respectively, and co-deliver them together with Alexa 647-HPV RNPs in live HeLa cells (Supplementary Figure 7**A**). 3-color imaging reveals robust and distinct puncta for Atto 488, TMR and Alexa 647, indicating efficient and specific simultaneous tagging of 3 separate genes. Higher multiplexing should also be achievable by adding more spectral channels and/or by spectral unmixing techniques.

### CAS-LiveFISH enables imaging spatial relations between HPV integration site and *MYC* in live HeLa cells

Proximity-ligation assays, such as Hi-C and ChIA-PET, suggested that activation of *MYC* in HeLa cells by the transcriptional enhancer created at the HPV integration site involves long-range genomic interactions (Figure 3**A**). However, the physical distances between enhancer and target gene at *MYC* have not been measured in live cells. To leverage CAS-LiveFISH for imaging spatial relationships between two genomic loci, we first create a pool of 288 *in vitro* transcribed gRNAs targeting ~40 kb around the *MYC* gene and assemble them with TMR-labelled dCas9 into fluorescent RNPs. Co-delivery of TMR-MYC and Alexa 647-HPV RNPs in live HeLa cells results in Alexa 647 puncta that often co-localize with TMR puncta (Figure 3**D**). As a control, co-delivery of TMR-Gal4 and Alexa 647-HPV RNPs results in Alexa 647 puncta but no discernible TMR puncta (Supplementary Figure 8**A**), further indicating specific tagging of the two genomic loci with minimal crosstalk.

To measure the distances between TMR-MYC and Alexa 647-HPV we first perform a two-color nanometer registration procedure using fluorescent nanoparticles, which achieves a root-mean-square (r.m.s.) registration accuracy of 21nm in *x*, 25nm in *y* and 32nm in 2D (Supplementary Figure 9**A**). We further validate the precision of distance measurements in live cells using dual TMR-HPV and Alexa 647-HPV tagging. The two-color HPV-HPV distances have a tight distribution with an average 2D distance of ~60nm, consistent with σ_*xy*_ ~39 nm 2D localization errors in each color (σ_*x,y*_ ~27 nm each in *x*, *y*) and 32 nm two-color 2D registration errors (Supplementary Figure 10**A-B**). Using this two-color nanometer co-localization procedure, we measure the relative distances between TMR-MYC and Alexa 647-HPV nuclear puncta. We observe a range of 2D MYC-HPV distances, with the majority (~75%) being within 200nm proximity, while a smaller sub-population (~10%) is further separated, at distances of 300nm-1μm (Figure 3**E**). These results illustrate how two-color tagging of genomic loci can provide insights into spatial relations of *cis*-elements and target genes, with nanometer resolution in live cells.

### Pre-assembled RNPs enable tracking the real-time motion of single gene loci

Understanding the movements of genomic loci in the nucleus is important for elucidating key processes such as gene regulation, DNA translocations and recombination. We use dCas9 RNP tagging and analyze the movement of the *MUC4* gene as well as the HPV integration site in live HeLa cells (Figure 4**A**). The mean-squared-displacement (MSD) vs. time of both *MUC4* and HPV follow a scaling behaviour, MSD~*Dt*^α^, with α<1 (Figure 4**B**), indicating sub-diffusive motion, typical for tagged genomic loci. For *MUC4* α≈0.5, in close agreement with previously observed α≈0.5 for other endogenous genes and *cis*-elements (Gu et al., 2018; Li et al., 2019). Scaling exponent α=0.5 characterizes the movement of individual monomers in a polymer chain (Rouse model) (Zhang and Dudko, 2016). Interestingly, the motion of HPV appears strongly sub-difussive, with α≈0.35, significantly deviating from the free polymer-like physical picture. This confined motion could reflect spatial constraints such as local compartmentalization, highly cross-linked chromatin chains or other possible mechanisms (Khanna et al., 2019; Zhang and Dudko, 2016). We also analyzed the velocity autocorrelation function (ACF) for the *MUC4* and HPV traces (Weber et al., 2010). The velocity ACFs for both loci exhibit negative dips (Figure 4**C**), indicating that the movement dynamics might be dominated by restoring forces often occurring inside a viscoelastic environment (Lucas et al., 2014; Weber et al., 2010; Zhang and Dudko, 2016). Both loci exhibit self-similar motions, as the velocity ACFs collapse upon time-axis rescaling. The HPV velocity ACF exhibits a more pronounced negative dip than *MUC4*, consistent with the more strongly confined nature of the HPV motion vs. *MUC4*. Taken together, these results demonstrate how pre-assembled fluorescent RNP tagging can provide quantitative insights into the detailed movements of specific genomic loci in the nucleus of live cells.

**Figure 4.**
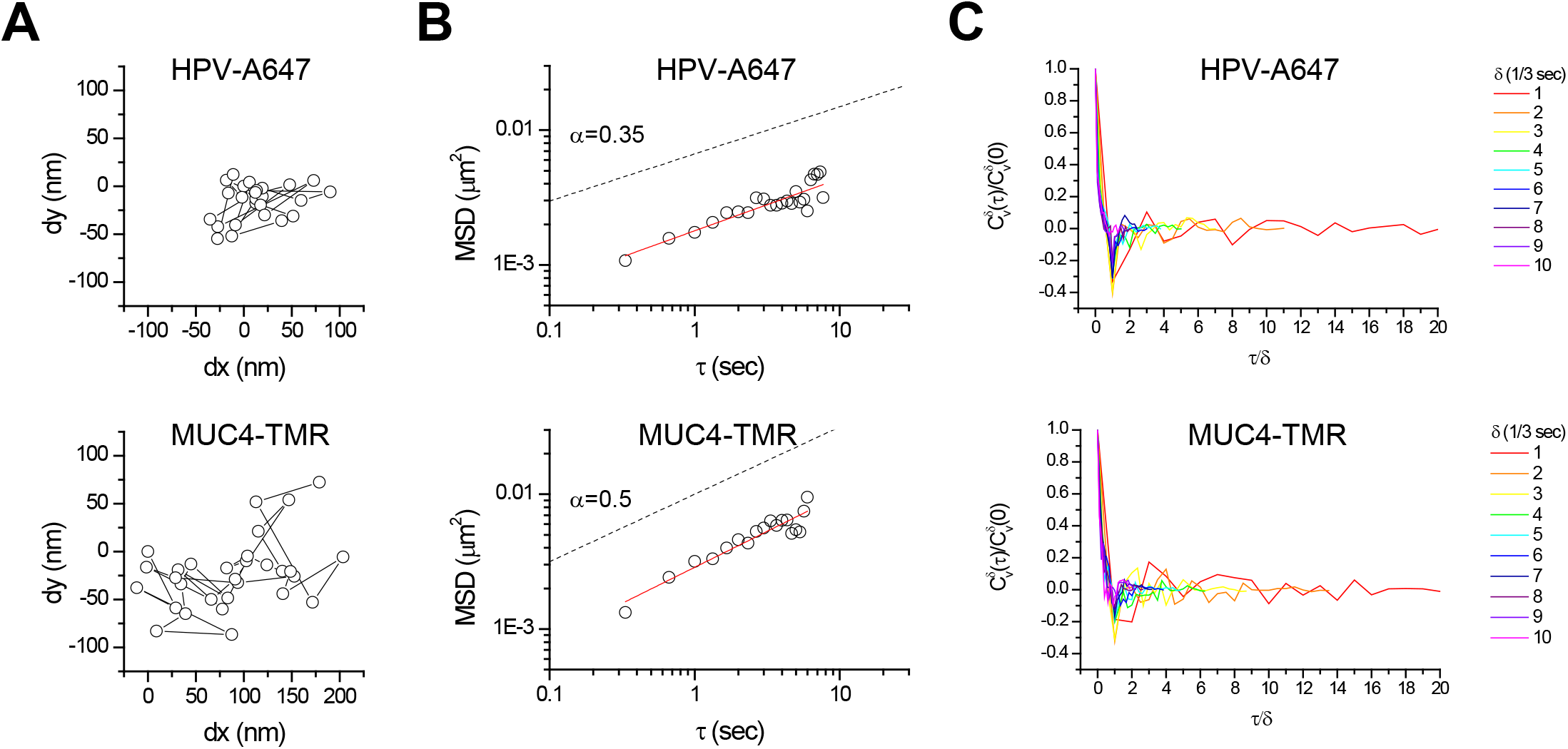
CAS-LiveFISH enables tracking the real-time motion of single gene loci. (**A, B, C**) Real-time tracking of Alexa 647-HPV (top) and TMR-MUC4 (bottom) motions. (**A**) Scatter plots showing the trajectory of single Alexa 647-HPV and TMR-MUC4 loci. (**B**) MSD vs. τ graphs show sub-diffusive motion, which is significantly stronger for HPV vs. *MUC4*. Red lines: linear fits, with α=0.37±0.04 and α=0.53±0.05 (*n*=31 and 15 loci) for HPV and *MUC4* respectively. (**C**) Velocity autocorrelation functions, showing negative dips, indicative of movement inside a viscoelastic environment. HPV exhibits more pronounced negative dips than *MUC4*, indicating stronger confinement. The time axis is rescaled with the time delay parameter δ; collapse of the curves indicates self-similar motions.

## DISCUSSION

Here we demonstrate CAS-LiveFISH, an approach for visualizing specific genomic loci in live mammalian cells and early embryos. CAS-LiveFISH relies on cell delivery of pre-assembled fluorescent dCas9-gRNA RNPs, bypassing the need for extensive genetic manipulations and optimization of multiple expression constructs.

Previous works had begun to recognize the versatility of pre-assembled RNPs for genomic imaging. A variant of FISH (CAS-FISH), based on fluorescent dCas9 RNPs instead of DNA hybridization probes (Deng et al., 2015), showed rapid, cost-effective and convenient multi-color labelling of genomic loci, in fixed cells. More recently, use of dCas9 RNPs with fluorescent CRISPR RNAs (crRNAs) achieved high SNR detection of highly-repetitive genomic loci in cell lines, as well as in primary human cells (Wang et al., 2019). The latter study also illustrated the potential of RNP-based approaches for medical diagnostics and for genomic imaging in systems that are not readily amenable to genetic manipulations. Here we significantly expand these previous works by demonstrating robust and efficient imaging of specific low-repeat containing and non-repetitive genomic loci, in live cells. dCas9 RNPs with fluorescent crRNAs could potentially achieve higher detection SNR, however use of fluorescent dCas9 and non-fluorescent *in vitro* transcribed gRNAs is significantly more cost-effective than pools of 10s-100s of individually chemically synthesized and purified fluorescent crRNAs. Future developments of low-cost methods to prepare *in vitro* transcribed pools of fluorescent gRNAs, such as using dye-binding aptamers (Autour et al., 2018; Filonov et al., 2014), could enable increased detection SNR, due to the rapid degradation of background gRNAs that are not “protected” in a target DNA-bound dCas9:RNA:DNA ternary complex. Finally, combining our DNA targeting RNP system with RNA targeting CRISPR proteins, such as dCas13 (Wang et al., 2019; Yang et al., 2019), would facilitate simultaneous imaging of *cis*-elements, target genes and nascent transcripts, as well as various other chromatin associated RNAs. Such direct visualization approaches would transform our ability to probe the links between genome organization and gene expression regulation as well as the nuclear organization, dynamics and function of non-coding regulatory RNAs.

CAS-LiveFISH by delivery of pre-assembled dCas9 RNPs can also be extended to imaging genomic loci in live early mouse embryos. The fluorescent RNPs are stable enough to potentially image over ~1 day-long time-scales. With emerging microscopy techniques to image in thick multi-cellular samples with increased sensitivity and resolution (Liu et al., 2018; McDole et al., 2018; Royer et al., 2016), imaging fainter signals, down to single non-repetitive genomic loci, and possibly together with simultaneous and coincidental single locus activity readouts and detection of regulatory molecules should also be within reach. These new capabilities could facilitate visualizing several important genomic processes at specific gene loci, in the context of early mammalian development.

A long-standing “mystery” of genome biology is how distal enhancers communicate with target genes to activate transcription (Furlong and Levine, 2018). Mechanisms involving direct physical interactions, such as long-range chromatin looping, are thought to facilitate this process, but it has been very challenging to directly measure the spatial relationships between distal enhancers and target genes in live cells. Progress has been made by introducing artificial exogenous tags, such as arrays of target sites for bacterial DNA-binding proteins (Alexander et al., 2019; Chen et al., 2018). CAS-LiveFISH enables directly tagging and imaging the endogenous genomic sequences, without needing genomic integrations of exogenous tags and possibly better reflecting the native chromatin organization. Our observations in live HeLa cells reveal that the distal enhancer created at the HPV integration site is often in <200nm proximity to the target *MYC* gene. An emerging view of distal gene regulation mechanisms proposes that approximate enhancer-promoter proximity (Alexander et al., 2019; Furlong and Levine, 2018), rather that direct molecular contact, might be adequate for transcription activation. High-local concentrations of Pol II regulatory factors (RFs) created around the enhancer might thus facilitate such action-at-a-distance. Interestingly, the physical distances are within the range (~200nm) over which Pol II RFs cluster in the vicinity of pluripotency genes like *Pou5f1* (aka *Oct4*) and *Nanog* in embryonic stem cells (Li et al., 2019). CAS-LiveFISH, combined with molecular imaging of Pol II RFs could further elucidate the physical nature and underlying genome topologies behind these important regulatory macromolecular assemblies.

Finally, the observations of highly sub-diffusive motion of the HPV integration site in HeLa cells are particularly striking. Previous work had also showed strongly sub-diffusive motions for Tetracycline operator arrays integrated in immunoglobulin gene segments in B-lymphocytes (Khanna et al., 2019). It was further postulated that such highly constrained motion might reflect a network of cross-linked chromatin, close to the boundary of a sol-gel phase transition. Intriguingly, such a picture might also apply to highly-active clusters of transcriptional enhancers, bound by high densities of transcription factors and various co-activator proteins (Furlong and Levine, 2018; Hnisz et al., 2017). The HPV-16 integration site in a related cervical neoplasia cell line colocalizes with a prominent nuclear focus that contains large amounts of Brd4, Mediator and H3K27 acetylated chromatin (Dooley et al., 2016). Further extensions to simultaneously visualize the HPV integration site in HeLa cells, together with RNA Polymerase II and various Pol II regulatory factors, as recently demonstrated for endogenous genes in mouse embryonic stem cells (Li et al., 2019), will be critical for testing various models for the physical organization of this important class of transcription regulatory units and their links to activation of important oncogenes like *MYC*.

## ACKNOWLEDGEMENTS

We thank Pai-Tseng Kuo and Hsin-Wei Wen (MSKCC Structural Biology) for assistance in creating the dCas9-GFP U2 O-S cell line, Vidur Garg and Kat Hadjantonakis (MSKCC Developmental Biology) and Willie Mark (MSKCC Mouse Genetics Core Facility) for assistance with mouse embryo microinjection, Luciano Marraffini (Rockefeller University) for useful discussions and Luke Lavis (HHMI/Janelia) for dye-labeling reagents. This work was supported by the Louis V. Gerstner, Jr. Young Investigators Fund (A.P.), a National Cancer Institute grant (P30 CA008748), a National Institutes of Health (NIH) Director’s New Innovator Award (1DP2GM105443-01; A.P.) and the National Institute of General Medical Sciences of NIH (1R01GM135545-01 and 1R21GM134342-01; A.P.).

## AUTHOR CONTRIBUTIONS

A.P. conceived, designed and supervised the study. Y.G. developed and validated the experimental systems. Y.G. and A.P. performed experiments and analyzed the data. A.P. wrote the manuscript.

## METHODS

### Mammalian and bacteria cell culture

HeLa and U-2 OS cells were grown in MEM (Life Technologies) and McCoy’s 5A Medium (ATCC) respectively, without phenol-red, and supplemented with 10% fetal bovine serum (FBS) 100U/mL penicillin/streptomycin, 1mM Sodium Pyruvate, 1× Non-Essential Aminoacids solution (NEAA) and 2mM L-alanyl-L-glutamine (GlutaMAX). All mammalian cells were grown in a 5% CO_2_ atmosphere at 37°C, in a humidified incubator. *Escherichia coli* were cultured in Luria-Bertani (LB) or 2×YT medium, in a 37°C shaking incubator. Required antibiotics were added to the medium at the following concentrations: ampicillin: 100 μg/ml; chloramphenicol: 34 μg/ml; kanamycin: 50 μg/ml. Bacterial cell growth was monitored periodically by determining the optical density of culture aliquots at 600 nm using NanoDrop 2000 (Thermo Scientific).

### Recombinant dCas9 expression and purification

The gene encoding a catalytically inactive *Streptococcus pyogenes* Cas9 (D10A/H840A) (dCas9) was cloned into a pD451-SR vector that contained a synthetic DNA fragment comprised of SNAPf or CLIPf tag, a Prescision protease site and a His10 affinity tag (synthesized by DNA2.0). The dCas9 sequence also contains two copies of a Nuclear Localization Signal (NLS) sequence, to facilitate nuclear import. The *S. pyogenes* Cas9 D10A/H840A mutant was expressed in *E. coli* strain BL21 Rosetta 2 (DE3) (Novagen) and cultured in 2×YT medium at 18°C for 16 h following induction with 0.2 mM IPTG. Cells were lysed in 20 mM Tris pH 8.0, 500 mM NaCl, 1 mM TCEP, supplemented with protease inhibitor cocktail (Roche). Clarified lysate was mixed with Ni-NTA agarose at 4℃ for 2h (Qiagen). Bound protein was eluted in 20 mM Tris pH 8.0, 250 mM NaCl, 10% glycerol and 150mM imidazole. The eluted protein were dialyzed overnight against 20 mM HEPES pH 7.5, 150 mM KCl, 1 mM TCEP, 10% glycerol and cleaved with PreScission protease to remove the His_10_ affinity tag. The cleaved dCas9 protein was further purified with a 5 ml SP Sepharose HiTrap column (GE Life Sciences) in 20 mM HEPES pH 7.5, 150 mM KCl, 1 mM TCEP, 10% glycerol with a linear gradient of 150 mM – 1 M KCl. Fractions containing dCas9 protein were pooled and the protein was concentrated with spin concentrators (Amicon Ultra 15, MWCO 10k; Millipore), and after adding glycerol to 20% final concentration, flash-frozen in liquid nitrogen and stored at −80°C.

### Fluorescent dCas9 labelling

For SNAP tag labelling, dCas9-SNAP protein samples were incubated with SNAP-Surface Alexa Fluor 647 (NEB S9136) at 37°C for 30 minutes. For CLIP tag labelling, CLIP-Cell TMR star substrate or CLIP-Surface 488 substrate (NEB S9219 and S9232) was reacted with dCas9-CLIP proteins at 37°C for 60min. The excess fluorescent SNAP or CLIP substrate was removed with three washes using 10 K molecular weight (MWCO) spin concentrators (Millipore). Fluorescent dCas9 protein was stored in 20 mM HEPES pH 7.5, 150 mM KCl, 1 mM TCEP, 10% glycerol. Protein concentration and labelling efficiency were determined by measuring absorption specta using a NanoDrop 2000 (Thermo Scientific). Calculated extinction coefficients at 280nm are 141,670 and 140,180 for dCas9-SNAP and dCas9-CLIP respectively. Fluorescent dCas9 protein was flash-frozen in liquid nitrogen and stored at −80°C.

### Preparation of *In vitro* transcribed gRNAs

A synthetic DNA fragment (IDT Gblocks) containing the full-length sequence of an optimized SP sgRNA (Chen et al., 2013) was inserted into the BamHI and EcoRI sites of a modified pLVX-shRNA2 vector (Clonetech) in which ZsGreen1 had been replaced by BFP. DNA templates for T7 transcription were prepared by PCR reactions using a common reverse primer and unique forward primers containing the T7 promoter and the seed sequence. The sgRNAs were *in vitro* synthesized with HiScribe™ T7 High Yield RNA Synthesis Kit (NEB). Transcription reactions were treated with RNase-free DNaseI (NEB) to digest the DNA template at 37°C for 1 hour. The sgRNAs were purified by phenol:chloroform: isoamyl alcohol (25:24:1) extraction and ethanol precipitation. The sgRNA pellets were dissolved in 20 mM HEPES (pH 7.5), 150 mM KCl, 10% glycerol and 1 mM TCEP. To refold purified sgRNAs, the sgRNAs were incubated at 70°C for 5min and slowly cooled down to room temperature. MgCl_2_ was then added to 1mM final concentration and the sgRNA samples were incubated at 50°C for 5min and slowly cooled down to room temperature. The sgRNAs concentration was estimated based on absorption measurements with a NanoDrop 2000 sprectrophotometer (Thermo Scientific) and the samples were stored at −80°C. The sequences of nucleic acids used in this study can be found in Supplementary Tables 1-3.

### Binding and Electrophoretic mobility shift assays

Fluorescent dCas9-gRNA RNP complexes were assembled by incubating fluorescent dCas9 protein with a ~30-fold molar excess of sgRNA for 10min at 37°C. DNA binding reactions contained 5 nM Cy3- or Cy5-labelled DNA duplex and increasing concentrations of dCas9-sgRNA RNPs. In competition assays, unlabelled DNA duplex was added at 5 nM. DNA binding reactions were incubated for 15min at 37°C. dCas9-sgRNA:DNA tertiary complex formation was analyzed by 4% polyacrylamide native gel electrophoresis in 0.5×TBE buffer. Fluorescent bands in the gels were visualized using a fluorescent scanner (Typhoon; GE).

### Delivery of sgRNAs and dCas9-gRNA RNPs into live cells and embryos

U-2 OS cells stably expressing spdCas9-GFP were nuclofected with Amaxa® Cell Line Nucleofector® Kit V (Lonza). Cells were detached with 0.05% trypsin/EDTA and spun down by centrifugation at 1000rpm for 5 min. 1×10^6^ cells were resuspended in 100 μl room-temperature Nucleofector® Solution and mixed with sgRNA and a LifeAct-BFP expressing plasmid. Each sample was transferred into a cuvette and nucleofected using Nucleofector® Program X-001. 500 μl of pre-equilibrated culture medium was then added to the cuvette and the cells were gently transferred into an 8-chamber coverglass (LabTek, 155411). Cells were imaged 4-6hr and 24 hrs after nucleofection.

To deliver fluorescent dCas9-gRNA RNP complexes into live cells, HeLa and U-2 OS cells were plated on 35mm cover-glass bottom dishes (MatTek) and microinjected with fluorescent dCas9-sgRNA RNP complexes. The fluorescent dCas9 RNP complexes were reconstituted by combining fluorescent dCas9 protein with sgRNA at a molar ratio of 1:30~1:50 in 20 mM HEPES (pH 7.5), 150 mM KCl, 10% glycerol and 1 mM TCEP at 37°C for 10min. Assembled RNP complexes were further diluted in microinjection buffer to about 100nM-1μM final concentration and filtered by spinning down at 2000 rpm for 2 min using a 0.22 μm sterilized centrifuge filter (COSTAR). The fluorescent dCas9-gRNA RNP complexes were loaded into glass microinjection capillary tips (Femtotips; Eppendorf) using a microloader pipette tip (Eppendorf) and injected into the nucleus of HeLa or U-2 OS cells using an Eppendorf FemtoJet® Microinjector (Eppendorf). After media were changed, the cells were left to recover in the 37°C/5% CO_2_ incubator for ~3-5 hours before imaging. For imaging genomic loci in mouse embryos, 2-cell stage mouse embryos were placed in FHM media under mineral oil in a round-bottom glass slide. After microinjection of fluorescent dCas9-gRNA RNPs, embryos were transferred to a 35mm cover-glass bottom dish, into drops of KSOM media under mineral oil, and placed in a 37°C/5% CO_2_ incubator until imaging.

### Live-cell and live-embryo imaging

Imaging was performed with a home-built microscope setup, featuring a 60× 1.49 NA objective lens (Nikon MRD01691), 405nm (Thorlabs diode), 488nm (Coherent Sapphire HP 500mW), 532nm (Coherent Verdi G2 2000mW) and 640nm (Coherent Cube 100mW) excitation lasers in epifluorescence configuration, a quad-view 4-color imaging device (Photometrics) and an EM-CCD camera (Ixon3 897; Andor). Imaging was performed at 3 frames/sec. Fine focusing was achieved by translating the sample in *z* with a nanopositioning stage (P-517.3CD, with E-710.3CD controller; Physik Instrumente). A home-built stage incubator and a separate objective lens heater maintained the sample at 37°C. The stage incubator atmosphere was adjusted with independent O_2_, N_2_ and CO_2_ mass-flow controllers (Omega). For some of the embryo imaging experiments, to achieve a larger field-of-view fitting whole embryos, we used a second epifluorescence microscope (Zeiss, Axiovert 200), equipped with an oil-immersion objective lens (Zeiss, Plan-apochromat 63 × 1.4 NA), a mercury lamp illuminator (VSG HBO100/001-26E), filter sets for DAPI (used for BFP), GFP, TMR (used for RFP) and Cy5 (used for SiR), and a scientific CCD (Hamamatsu, ORCA Flash part #C47428012AG).

### Image and data analysis

Image analysis was performed in IDL (ITT Visual Information Solution, version 8.0.1). First, nuclear puncta were identified using as local maxima in band-pass filtered and background-subtracted images (Crocker and Grier, 1996). Then, to obtain more precise *xy* coordinates, the processed images were fitted to a 2D elliptical Gaussian peak function using non-linear least-squares fitting. For calibrating the registration between Alexa 647 and TMR images, we used the coordinates of 100nm Tetraspek beads (ThermoFisher, T7284) scattered throughout the field of view. Two-color registration (Pertsinidis et al., 2010) was performed using a second-order polynomial spatial warping transformation (Pertsinidis et al., 2013; Wang et al., 2016). MSD and velocity ACFs were calculated with a MATLAB routine (Matworks, version 2010b). Observed MSDs were corrected by subtracting a constant, MSD^loc_error^, to account for the effect of localization errors (Kepten et al., 2013). MSD^loc_error^ was estimated by tracking the relative motion of a single genomic locus, simultaneously tagged with dCas9-Alexa 647 and dCas9-TMR (Khanna et al., 2019). Graphing, linear least-squares fitting and statistical analysis was performed using Origin (OriginLab, version 8.5.0).

