## Supplementary Information for "CAS-LiveFISH: Simple and versatile imaging of genomic loci in live mammalian cells and early pre-implantation embryos"

[2] *Lead Contact*

#### Supplementary Information

1. **Supplementary Figures 1-9**
2. **Supplementary Material**

    DNA Sequences

3. **Supplementary Tables 1-3**

#### Supplementary Figures

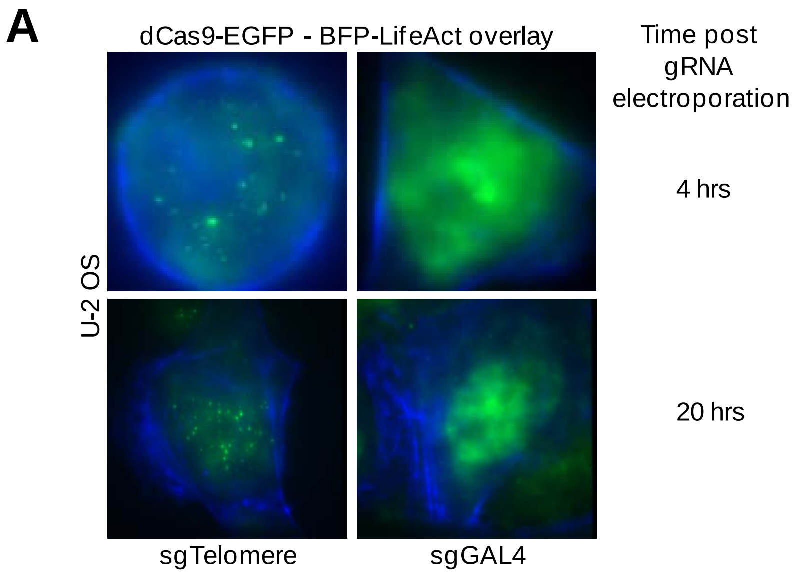


**Supplementary Figure 1. Imaging of genomic loci by delivery of *in vitro* transcribed gRNAs.** (**A**) Visualization of telomeres in U-2 OS cells stably expressing dCas9-EGFP, by electroporation. Examples are shown at 4 hrs and 20 hrs after electroporation. No nuclear puncta are observed for an sgGAL4 non-targeting control.

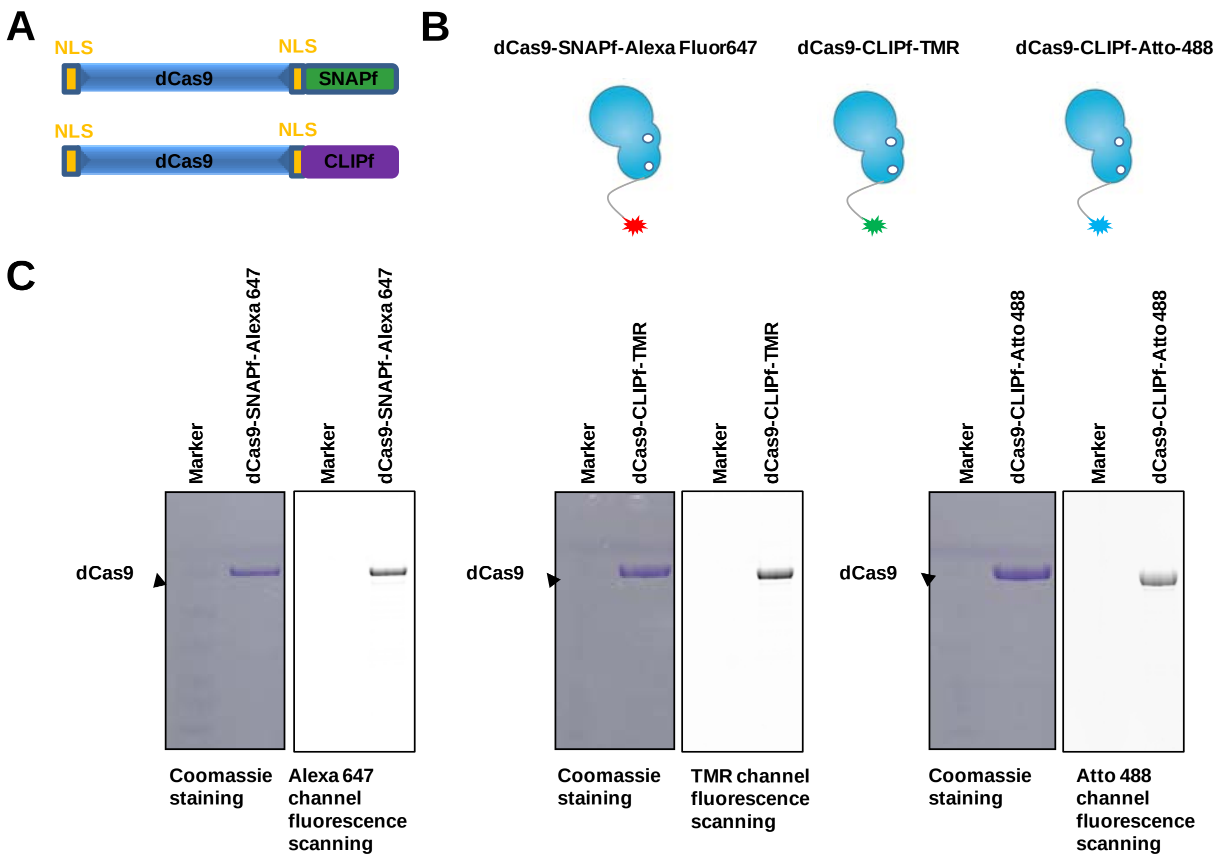


**Supplementary Figure 2. Fluorescent dCas9 proteins.** (**A, B**) Schematic of dCas9-SNAPf and dCas9-CLIPf constructs. (**C**) SDS-PAGE showing purified and fluorescently labelled dCas9 proteins.

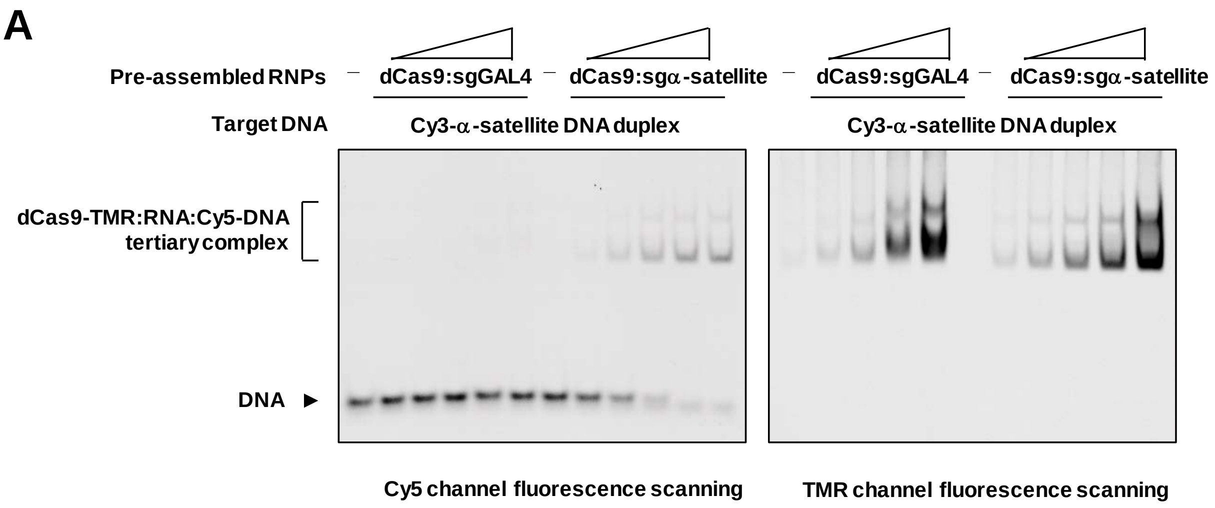


**Supplementary Figure 3. Electrophoretic mobility shift assays.** (**A**) Pre-assembled dCas9-TMR RNPs were incubated with Cy5-DNA duplexes and ternary complexes are resolved from free DNA using native SDS-PAGE. Binding is observed for the targeting gRNA (sgα-satellite) but not for the non-targeting control (sgGAL4).

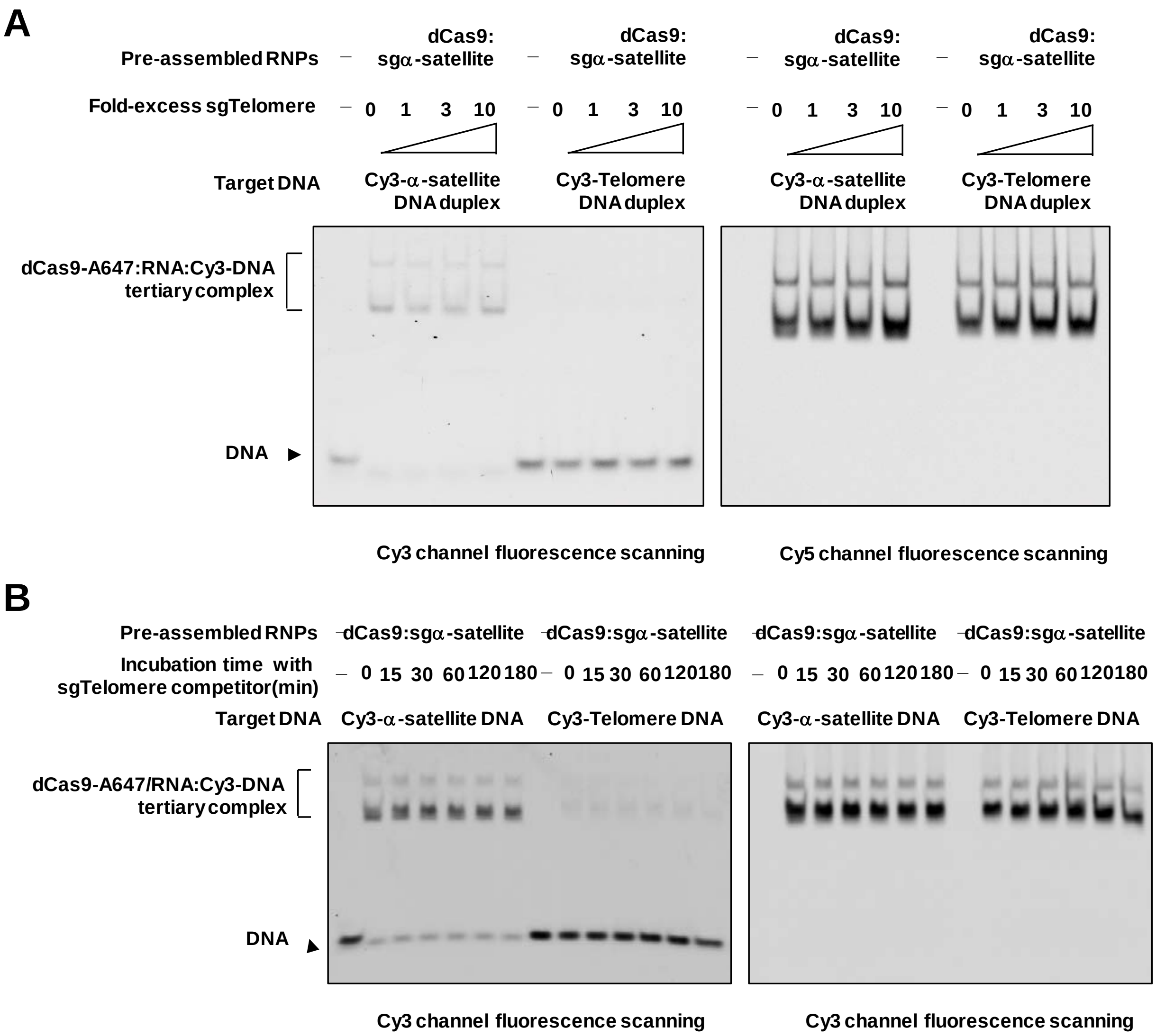


**Supplementary Figure 4. Biochemical characterization of gRNA cross-talk/competition.** (**A**) dCas9-Alexa 647 RNPs are assembled with a gRNA against α-satellite sequences (sgα-satellite) and incubated with 1-10-fold excess of a competitor gRNA (sgTelomere). Binding is only observed for the Cy3-α-satellite DNA target duplex but not for the Cy3-Telomere DNA duplex. (**B**) Time course of competition experiment. Very little binding to the Cy3-Telomere DNA duplex is observed, even after 180 minute-long incubation.

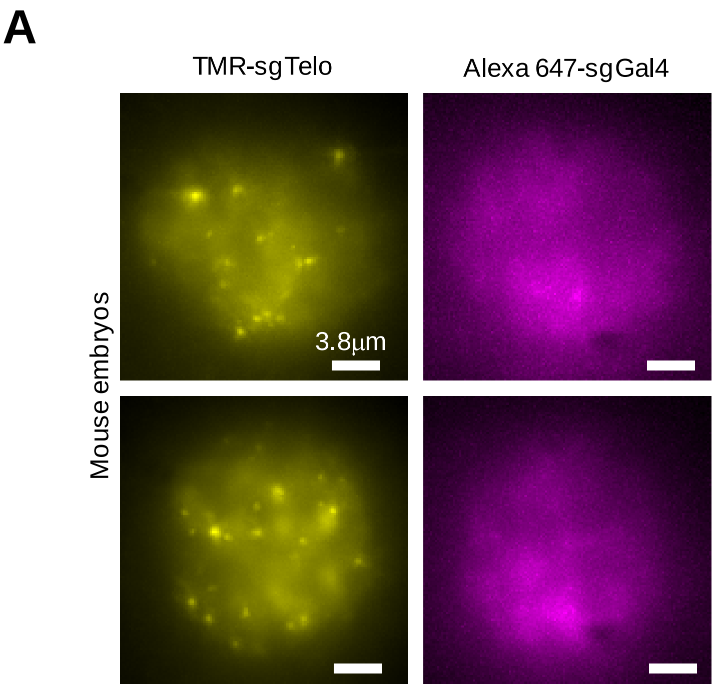


**Supplementary Figure 5. Specificity of fluorescent dCas9-gRNA RNPs delivered in live 2-cell-stage mouse embryos.** (**A**) Nuclear puncta are observed for dCas9-TMR assembled with a targeting gRNA (sgTelo), but no puncta are observed for dCas9-Alexa 647 assembled with a non-targeting gRNA (sgGal4).

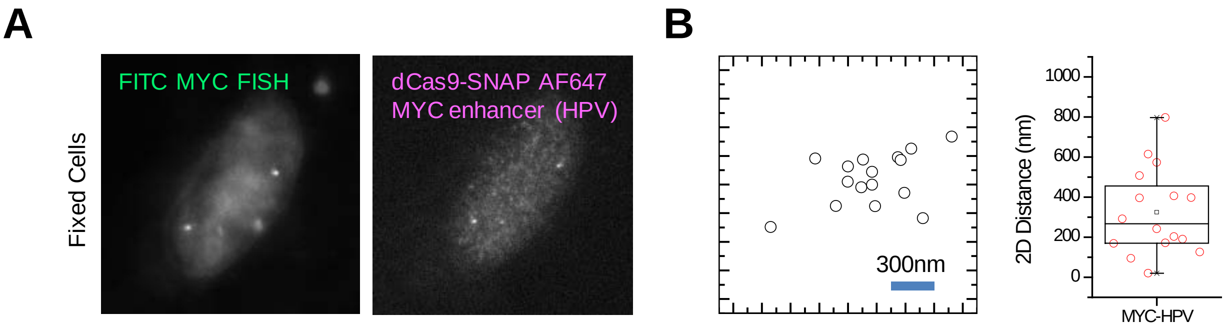


**Supplementary Figure 6. Validation of dCas9-HPV tagging using DNA FISH in fixed HeLa cells.** (**A**) Two-color imaging of FITC MYC FISH probes and dCas9-Alexa 647-HPV RNPs. (**B**) Scatter plot and (**C**) 2D distances between HPV loci vs. the centroid of the *MYC* FISH probes.

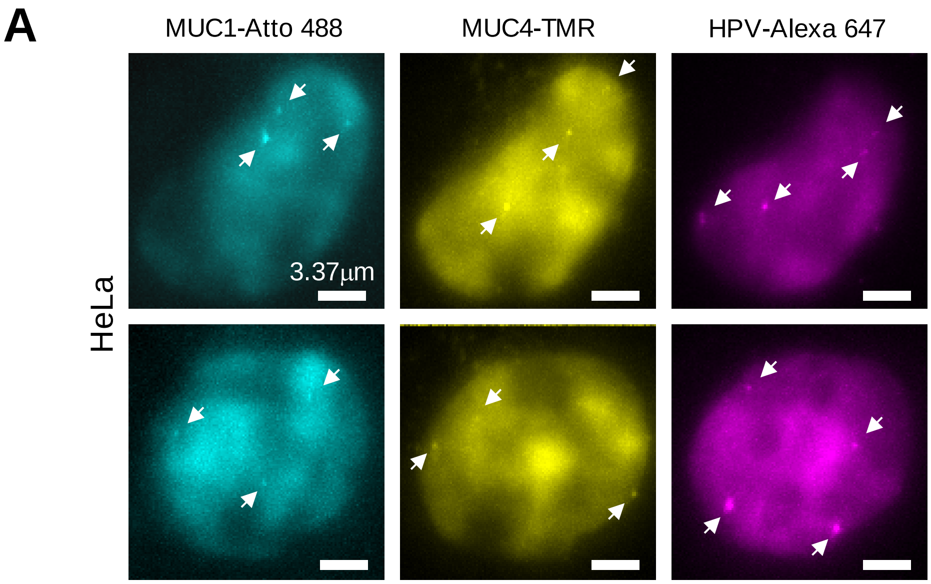


**Supplementary Figure 7. Multiplex imaging of 3 genes.** (**A**) Atto 488-MUC1, TMR-MUC4 and Alexa 647-HPV RNPs co-delivered in live HeLa cells. Arrows show nuclear puncta.

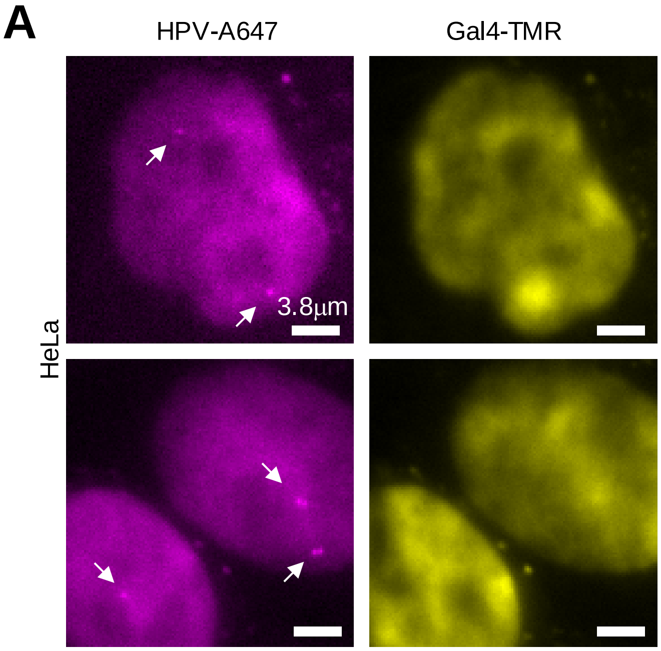


**Supplementary Figure 8. Control experiments for two-color imaging of genomic loci in live HeLa cells.** (**A**) Imaging of co-delivered Alexa 647-HPV and Gal4-TMR RNPs. Arrows indicate nuclear puncta for Alexa 647-HPV. No discernible nuclear puncta are seen for Gal4-TMR RNPs.

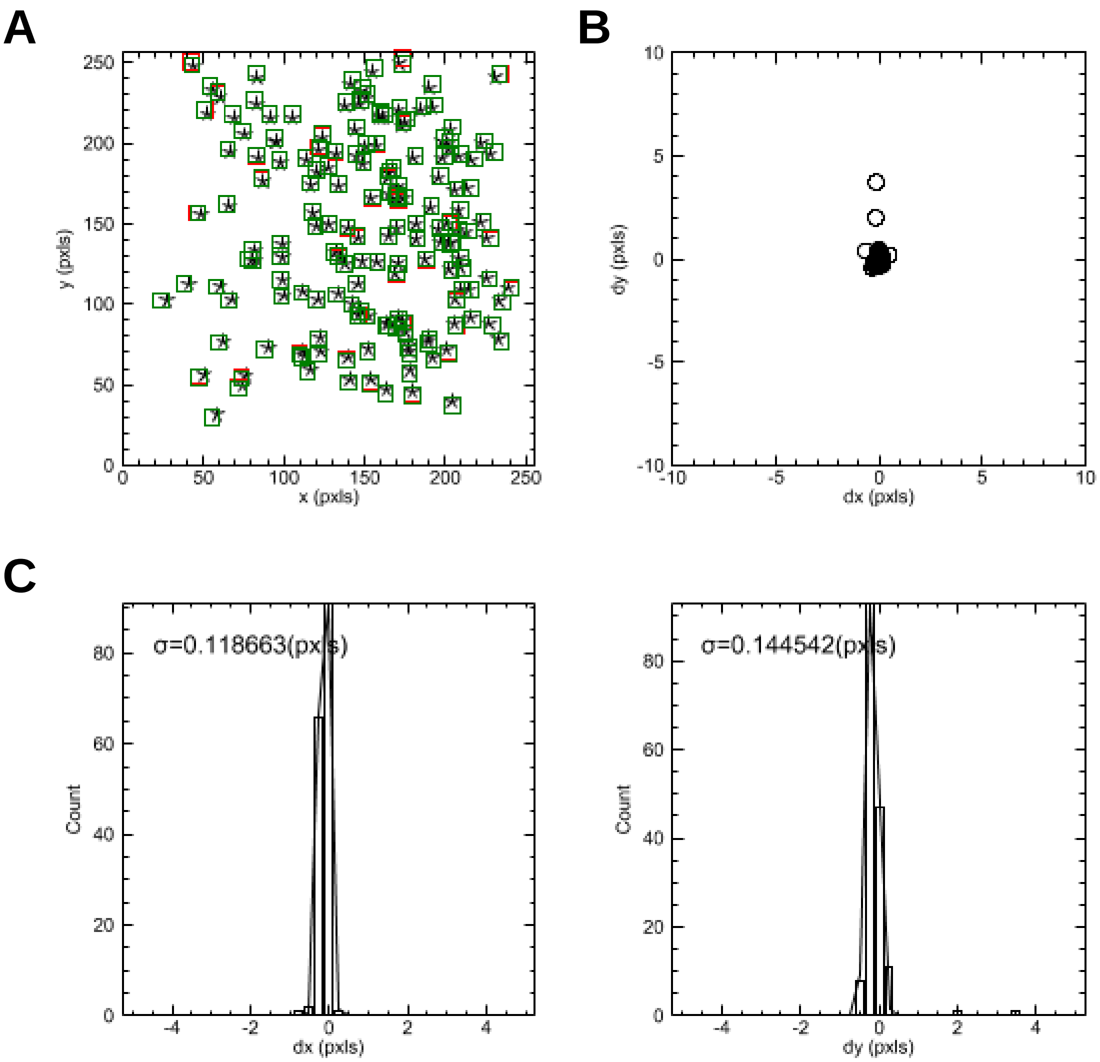


**Supplementary Figure 9. Two-color registration.** (**A**) Scatter plot of coordinates of 100 nm Tetraspek beads. (**B**) Scatter plot and (**C,D**) histograms of relative coordinates between TMR and Alexa 647 channel.

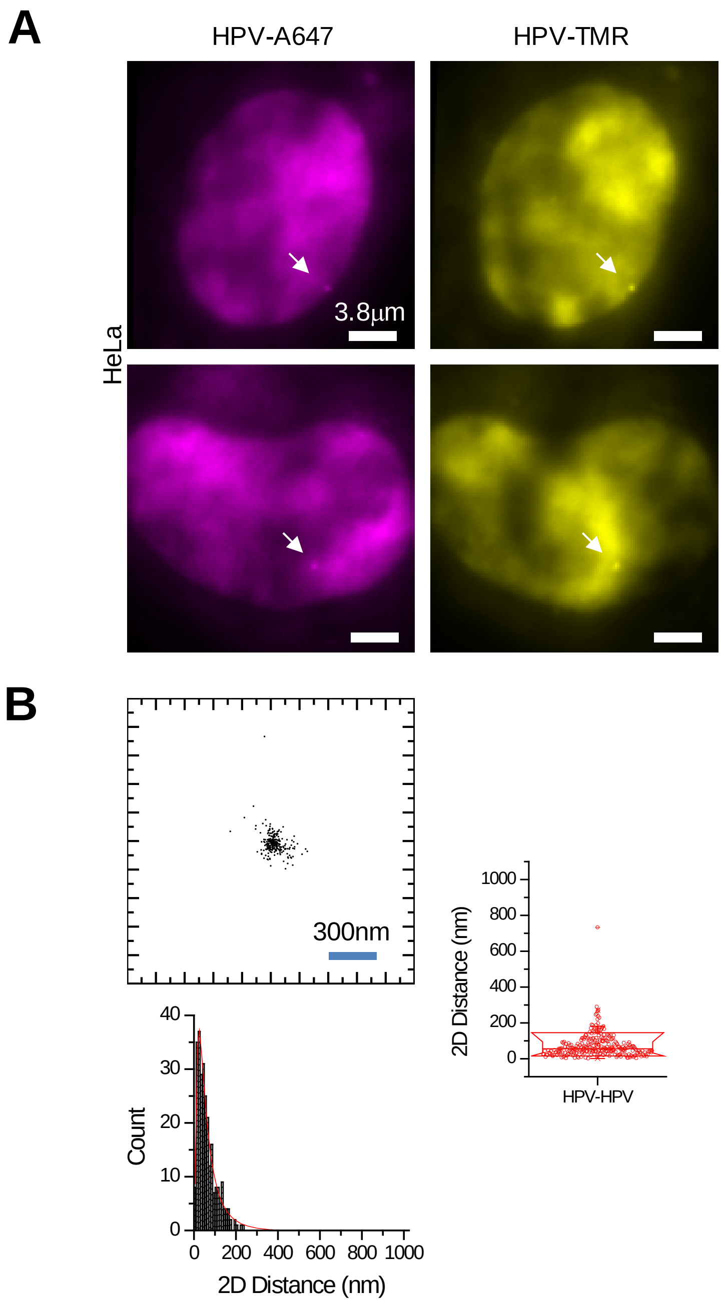


**Supplementary Figure 10. Characterization of two-color 2D distance measurement precision in live cells.** (**A**) Two-color imaging of HPV integration site, simultaneously in two colors (Alexa 647 and TMR) using co-delivery of preassembled RNPs in live HeLa cells. (**B**) 2D scatter plot of relative positions and statistics (histogram and box plot) of 2D distances between Alexa 647-HPV and TMR-HPV. 2D distance is 61±45 nm (mean±S.D.; *n*=271).

### Supplementary Material

#### DNA Sequences

##### Sequence of the dCas9-SNAP/CLIP Proteins

dCas9-SNAP fusion protein:

AGATCT*CCCAAAAAGAAGAGGAAAGTG***ATGGATAAGAAATACTCAATAGGCTTAGC TATCGGCACAAATAGCGTCGGATGGGCGGTGATCACTGATGAATATAAGGTTC CGTCTAAAAAGTTCAAGGTTCTGGGAAATACAGACCGCCACAGTATCAAAAAA AATCTTATAGGGGCTCTTTTATTTGACAGTGGAGAGACAGCGGAAGCGACTCG TCTCAAACGGACAGCTCGTAGAAGGTATACACGTCGGAAGAATCGTATTTGTTA TCTACAGGAGATTTTTTCAAATGAGATGGCGAAAGTAGATGATAGTTTCTTTCA TCGACTTGAAGAGTCTTTTTTGGTGGAAGAAGACAAGAAGCATGAACGTCATC CTATTTTTGGAAATATAGTAGATGAAGTTGCTTATCATGAGAAATATCCAACTA TCTATCATCTGCGAAAAAAATTGGTAGATTCTACTGATAAAGCGGATTTGCGCT TAATCTATTTGGCCTTAGCGCATATGATTAAGTTTCGTGGTCATTTTTTGATTGA GGGAGATTTAAATCCTGATAATAGTGATGTGGACAAACTATTTATCCAGTTGGT ACAAACCTACAATCAATTATTTGAAGAAAACCCTATTAACGCAAGTGGAGTAGA TGCTAAAGCGATTCTTTCTGCACGATTGAGTAAATCAAGACGATTAGAAAATCT CATTGCTCAGCTCCCCGGTGAGAAGAAAAATGGCTTATTTGGGAATCTCATTGC TTTGTCATTGGGTTTGACCCCTAATTTTAAATCAAATTTTGATTTGGCAGAAGA TGCTAAATTACAGCTTTCAAAAGATACTTACGATGATGATTTAGATAATTTATT GGCGCAAATTGGAGATCAATATGCTGATTTGTTTTTGGCAGCTAAGAATTTATC AGATGCTATTTTACTTTCAGATATCCTAAGAGTAAATACTGAAATAACTAAGGC TCCCCTATCAGCTTCAATGATTAAACGCTACGATGAACATCATCAAGACTTGAC TCTTTTAAAAGCTTTAGTTCGACAACAACTTCCAGAAAAGTATAAAGAAATCTT TTTTGATCAATCAAAAAACGGATATGCAGGTTATATTGATGGGGGAGCTAGCCA AGAAGAATTTTATAAATTTATCAAACCAATTTTAGAAAAAATGGATGGTACTGA GGAATTATTGGTGAAACTAAATCGTGAAGATTTGCTGCGCAAGCAACGGACCT TTGACAACGGCTCTATTCCCCATCAAATTCACTTGGGTGAGCTGCATGCTATTT TGAGAAGACAAGAAGACTTTTATCCATTTTTAAAAGACAATCGTGAGAAGATTG**

AAAAAATCTTGACTTTTCGAATTCCTTATTATGTTGGTCCATTGGCGCGTGGCA

ATAGTCGTTTTGCATGGATGACTCGGAAGTCTGAAGAAACAATTACCCCATGGA

ATTTTGAAGAAGTTGTCGATAAAGGTGCTTCAGCTCAATCATTTATTGAACGCA

TGACAAACTTTGATAAAAATCTTCCAAATGAAAAAGTACTACCAAAACATAGTT

TGCTTTATGAGTATTTTACGGTTTATAACGAATTGACAAAGGTCAAATATGTTA

CTGAAGGAATGCGAAAACCAGCATTTCTTTCAGGTGAACAGAAGAAAGCCATT

GTTGATTTACTCTTCAAAACAAATCGAAAAGTAACCGTTAAGCAATTAAAAGAA

GATTATTTCAAAAAAATAGAATGTTTTGATAGTGTTGAAATTTCAGGAGTTGAA

GATAGATTTAATGCTTCATTAGGTACCTACCATGATTTGCTAAAAATTATTAAA

GATAAAGATTTTTTGGATAATGAAGAAAATGAAGATATCTTAGAGGATATTGTT

TTAACATTGACCTTATTTGAAGATAGGGAGATGATTGAGGAAAGACTTAAAACA

TATGCTCACCTCTTTGATGATAAGGTGATGAAACAGCTTAAACGTCGCCGTTAT

ACTGGTTGGGGACGTTTGTCTCGAAAATTGATTAATGGTATTAGGGATAAGCAA

TCTGGCAAAACAATATTAGATTTTTTGAAATCAGATGGTTTTGCCAATCGCAAT

TTTATGCAGCTGATCCATGATGATAGTTTGACATTTAAAGAAGACATTCAAAAA

GCACAAGTGTCTGGACAAGGCGATAGTTTACATGAACATATTGCAAATTTAGCT

GGTAGCCCTGCTATTAAAAAAGGTATTTTACAGACTGTAAAAGTTGTTGATGAA

TTGGTCAAAGTAATGGGGCGGCATAAGCCAGAAAATATCGTTATTGAAATGGC

ACGTGAAAATCAGACAACTCAAAAGGGCCAGAAAAATTCGCGAGAGCGTATGA

AACGAATCGAAGAAGGTATCAAAGAATTAGGAAGTCAGATTCTTAAAGAGCAT

CCTGTTGAAAATACTCAATTGCAAAATGAAAAGCTCTATCTCTATTATCTCCAA

AATGGAAGAGACATGTATGTGGACCAAGAATTAGATATTAATCGTTTAAGTGAT

TATGATGTCGATGCCATTGTTCCACAAAGTTTCCTTAAAGACGATTCAATAGAC

AATAAGGTCTTAACGCGTTCTGATAAAAATCGTGGTAAATCGGATAACGTTCCA

AGTGAAGAAGTAGTCAAAAAGATGAAAAACTATTGGAGACAACTTCTAAACGC

CAAGTTAATCACTCAACGTAAGTTTGATAATTTAACGAAAGCTGAACGTGGAGG

TTTGAGTGAACTTGATAAAGCTGGTTTTATCAAACGCCAATTGGTTGAAACTCG

CCAAATCACTAAGCATGTGGCACAAATTTTGGATAGTCGCATGAATACTAAATA

CGATGAAAATGATAAACTTATTCGAGAGGTTAAAGTGATTACCTTAAAATCTAA

ATTAGTTTCTGACTTCCGAAAAGATTTCCAATTCTATAAAGTACGTGAGATTAA

CAATTACCATCATGCCCATGATGCGTATCTAAATGCCGTCGTTGGAACTGCTTT

**GATTAAGAAATATCCAAAACTTGAATCGGAGTTTGTCTATGGTGATTATAAAGT**
**TTATGATGTTCGTAAAATGATTGCTAAGTCTGAGCAAGAAATAGGCAAAGCAAC**
**CGCAAAATATTTCTTTTACTCTAATATCATGAACTTCTTCAAAACAGAAATTACA**
**CTTGCAAATGGAGAGATTCGCAAACGCCCTCTAATCGAAACTAATGGGGAAAC**
**TGGAGAAATTGTCTGGGATAAAGGGCGAGATTTTGCCACAGTGCGCAAAGTAT**
**TGTCCATGCCCCAAGTCAATATTGTCAAGAAAACAGAAGTACAGACAGGCGGA**
**TTCTCCAAGGAGTCAATTTTACCAAAAAGAAATTCGGACAAGCTTATTGCTCGT**
**AAAAAAGACTGGGATCCAAAAAAATATGGTGGTTTTGATAGTCCAACGGTAGC**
**TTATTCAGTCCTAGTGGTTGCTAAGGTGGAAAAAGGGAAATCGAAGAAGTTAA**
**AATCCGTTAAAGAGTTACTAGGGATCACAATTATGGAAAGAAGTTCCTTTGAAA**
**AAAATCCGATTGACTTTTTAGAAGCTAAAGGATATAAGGAAGTTAAAAAAGACT**
**TAATCATTAAACTACCTAAATATAGTCTTTTTGAGTTAGAAAACGGTCGTAAAC**
**GGATGCTGGCTAGTGCCGGAGAATTACAAAAAGGAAATGAGCTGGCTCTGCCA**
**AGCAAATATGTGAATTTTTTATATTTAGCTAGTCATTATGAAAAGTTGAAGGGT**
**AGTCCAGAAGATAACGAACAAAAACAATTGTTTGTGGAGCAGCATAAGCATTAT**
**TTAGATGAGATTATTGAGCAAATCAGTGAATTTTCTAAGCGTGTTATTTTAGCA**
**GATGCCAATTTAGATAAAGTTCTTAGTGCATATAACAAACATAGAGACAAACCA**
**ATACGTGAACAAGCAGAAAATATTATTCATTTATTTACGTTGACGAATCTTGGA**
**GCTCCCGCTGCTTTTAAATATTTTGATACAACAATTGATCGTAAACGATATACG**
**TCTACAAAAGAAGTTTTAGATGCCACTCTTATCCATCAATCCATCACTGGTCTT**
**TATGAAACACGCATTGATTTGAGTCAGCTAGGAGGTGAC**GAGGGAGCT*CCCAAG*
*AAAAAGCGCAAGGTA*GTCGACGGTGGTTCTGGTATGGACAAAGACTGCGAAATGAAG
CGCACCACCCTGGATAGCCCTCTGGGCAAGCTGGAACTGTCTGGGTGCGAACAGGG
CCTGCACCGTATCATCTTCCTGGGCAAAGGAACATCTGCCGCCGACGCCGTGGAAGT
GCCTGCCCCAGCCGCCGTGCTGGGCGGACCAGAGCCACTGATGCAGGCCACCGCCT
GGCTCAACGCCTACTTTCACCAGCCTGAGGCCATCGAGGAGTTCCCTGTGCCAGCCC
TGCACCACCCAGTGTTCCAGCAGGAGAGCTTTACCCGCCAGGTGCTGTGGAAACTGC
TGAAAGTGGTGAAGTTCGGAGAGGTCATCAGCTACAGCCACCTGGCCGCCCTGGCC
GGCAATCCCGCCGCCACCGCCGCCGTGAAAACCGCCCTGAGCGGAAATCCCGTGCC
CATTCTGATCCCCTGCCACCGGGTGGTGCAGGGCGACCTGGACGTGGGGGGCTACG
AGGGCGGGCTCGCCGTGAAAGAGTGGCTGCTGGCCCACGAGGGCCACAGACTGGGC

AAGCCTGGGCTGGGTCTCGAGGGTGGTTCTGGTCTGGAAGTTCTGTTCCAGGGGCCC
CATCATCACCATCACCACCATCATCACCATTAA

The CLIP sequence in pD451-SR-dCas9-CLIP plasmid is:

ATGGACAAAGACTGCGAAATGAAGCGCACCACCCTGGATAGCCCTCTGGGCAAGCT
GGAACTGTCTGGGTGCGAACAGGGCCTGCACCGTATCATCTTCCTGGGCAAAGGAA
CATCTGCCGCCGACGCCGTGGAAGTGCCTGCCCCAGCCGCCGTGCTGGGCGGACCA
GAGCCACTGATCCAGGCCACCGCCTGGCTCAACGCCTACTTTCACCAGCCTGAGGCC
ATCGAGGAGTTCCCTGTGCCAGCCCTGCACCACCCAGTGTTCCAGCAGGAGAGCTTT
ACCCGCCAGGTGCTGTGGAAACTGCTGAAAGTGGTGAAGTTCGGAGAGGTCATCAG
CGAGAGCCACCTGGCCGCCCTGGTGGGCAATCCCGCCGCCACCGCCGCCGTGAACA
CCGCCCTGGACGGAAATCCCGTGCCCATTCTGATCCCCTGCCACCGGGTGGTGCAGG
GCGACAGCGACGTGGGGCCCTACCTGGGCGGGCTCGCCGTGAAAGAGTGGCTGCTG
GCCCACGAGGGCCACAGACTGGGCAAGCCTGGGCTGGGT

Notes: *italics* indicate nuclear localization signal (NLS) sequence; **bold** text is the sequence of dCas9; underlined text is the SNAPf sequence; red text is GGSG linker sequence; blue text is HRV 3C site and purple text is $His_{10}$ affinity tag.

**Synthetic sgTelomere DNA for cloning into pLVX-BFP plasmid:**

ggatcc**GTTAGGGTTAGGGTTAGGGTTA**GTTTAAGAGCTATGCTGGAAACAGCATAG
CAAGTTTAAATAAGGCTAGTCCGTTATCAACTTGAAAAAGTGGCACCGAGTCGGTGC
TTTTTTTacgcgtgaattc

Note: bold text is the sequence of sgTelomere and underlined text is the sgRNA scaffold.

#### Supplementary Tables

##### Supplementary Table 1. Primers for *in vitro* T7 transcribed sgGAL4, sgTelomere, sgα-satellite, sgMUC4 and sgMUC1

| Primer name | Sequence 5’-3’ |
|---|---|
| Common REV | AAAAAAAGCACCGACTCGGTGCCAC |
| sgGAL4 FWD | GAAAT**TAATACGACTCACTATAG**GTTGGAGCACTGTCCTCCGAACGTGTTTAAG |
| sgTelomere FWD | GAAAT**TAATACGACTCACTATAG**GTTAGGGTTAGGGTTAGGGTTAG |
| sgα-satellite FWD | GAAAT**TAATACGACTCACTATAG**GTAGAATCTGCAAGTGGATATTGTT |
| sgMUC4 E3 FWD | GAAAT**TAATACGACTCACTATAG**GTGGCGTGACCTGTGGATGCTGGTTTAAGAGCTATGCTGGAAACA |
| sgMUC1 E3 FWD | GAAAT**TAATACGACTCACTATAG**GTCCGGGGCCGAGGTGACACCGTGTTTAAGAGCTATGCTGGAAACA |

**Bold**, T7 promoter； underlined, sequence of sgRNA target; FWD, forward primer; REV: reverse primer.

##### Supplementary Table 2. Primers for *in vitro* T7 transcribed gRNAs targeting the HPV-18 integration site in HeLa cells

| Primer name | Sequence 5’-3’ |
|---|---|
| sgHPV FWD1 | GAAAT**TAATACGACTCACTATAG**ATATTTGTCAAATGCCAAATGTTTAAGAGCTATGCTGGAAACA |
| sgHPV FWD2 | GAAAT**TAATACGACTCACTATAG**GAATGTTTAA |

| | |
|---|---|
| | CTTCTAGGCCGTTTAAGAGCTATGCTGGAAACA |
| sgHPV FWD3 | GAAAT**TAATACGACTCACTATAG**GAATGCCTTCAAAGAACAGCAGTTTAAGAGCTATGCTGGAAACA |
| sgHPV FWD4 | GAAAT**TAATACGACTCACTATAG**GCAATGCTTAACACGGCAGCGTTTAAGAGCTATGCTGGAAACA |
| sgHPV FWD5 | GAAAT**TAATACGACTCACTATAG**GCATTGCTAATCTAGAAGAAGTTTAAGAGCTATGCTGGAAACA |
| sgHPV FWD6 | GAAAT**TAATACGACTCACTATAG**GAAAAGAAGCAGAGGTATGCGTGTTTAAGAGCTATGCTGGAAACA |
| sgHPV FWD7 | GAAAT**TAATACGACTCACTATAG**GTTCAATTGGTATGATTTAAGTTTAAGAGCTATGCTGGAAACA |
| sgHPV FWD8 | GAAAT**TAATACGACTCACTATAG**ATAAGGGCCACATAATGGAGGTTTAAGAGCTATGCTGGAAACA |
| sgHPV FWD9 | GAAAT**TAATACGACTCACTATAG**GACATTAATCTTAAGTATCCAGTTTAAGAGCTATGCTGGAAACA |
| sgHPV FWD10 | GAAAT**TAATACGACTCACTATAG**GTGTGATGGGAGCCATGTGGTTTAAGAGCTATGCTGGAAACA |
| sgHPV FWD11 | GAAAT**TAATACGACTCACTATAG**GGCAACTTCAGGCCATAGTCACGTTTAAGAGCTATGCTGGAAACA |
| sgHPV FWD12 | GAAAT**TAATACGACTCACTATAG**GCTAGACTCATTCATGCATTCGTTTAAGAGCTATGCTGGAAACA |
| sgHPV FWD13 | GAAAT**TAATACGACTCACTATAG**GTTCAATCATTTAGAGAAGAGTTTAAGAGCTATGCTGGAAACA |

| | |
|---|---|
| sgHPV FWD14 | GAAAT**TAATACGACTCACTATAG**AAGGACTTCTTCATGTACCCGTTTAAGAGCTATGCTGGAAACA |
| sgHPV FWD15 | GAAAT**TAATACGACTCACTATAG**ATACACCGAGGGAAAAACCAGTTTAAGAGCTATGCTGGAAACA |
| sgHPV FWD16 | GAAAT**TAATACGACTCACTATAG**GTACCTGGCCAAGATTAATTGTTTAAGAGCTATGCTGGAAACA |
| sgHPV FWD17 | GAAAT**TAATACGACTCACTATAG**GTCTGTGCAAAAAAAGTGCAGGTTTAAGAGCTATGCTGGAAACA |
| sgHPV FWD18 | GAAAT**TAATACGACTCACTATAG**GAAGTTCTCCTCCAAGGAAGGTTTAAGAGCTATGCTGGAAACA |
| sgHPV FWD19 | GAAAT**TAATACGACTCACTATAG**GGATAATTTGGGTCTCAAAAGTTTAAGAGCTATGCTGGAAACA |
| sgHPV FWD20 | GAAAT**TAATACGACTCACTATAG**GCTTCTGGGGAAGAGGAGTAGTTTAAGAGCTATGCTGGAAACA |
| sgHPV FWD21 | GAAAT**TAATACGACTCACTATAG**ATAAGATAGGGACCTAGACAGTTTAAGAGCTATGCTGGAAACA |
| sgHPV FWD22 | GAAAT**TAATACGACTCACTATAG**GAGAAATGTTATTGAAACGTTTAAGAGCTATGCTGGAAACA |
| sgHPV FWD23 | GAAAT**TAATACGACTCACTATAG**GTGCAGAGTGACACAGTATTGTTTAAGAGCTATGCTGGAAACA |
| sgHPV FWD24 | GAAAT**TAATACGACTCACTATAG**GATGCCATTATAGACAAGAACGTTTAAGAGCTATGCTGGAAACA |

| | |
|---|---|
| sgHPV FWD25 | GAAAT**TAATACGACTCACTATAG**GTAGAACAGACAGGACTACAGTTTAAGAGCTATGCTGGAAACA |
| sgHPV FWD26 | GAAAT**TAATACGACTCACTATAG**GCCAACTAGCCTCAAATTAAAGCAGTTTAAGAGCTATGCTGGAAACA |
| sgHPV FWD27 | GAAAT**TAATACGACTCACTATAG**GCCCCAAATCCAGGTTCCCCGTTTAAGAGCTATGCTGGAAACA |
| sgHPV FWD28 | GAAAT**TAATACGACTCACTATAG**GCTTACGGATCTACTTCTAATGTTTAAGAGCTATGCTGGAAACA |
| sgHPV FWD29 | GAAAT**TAATACGACTCACTATAG**GCTCCACCAAGATGGGAAGGAGTTTAAGAGCTATGCTGGAAACA |
| sgHPV FWD30 | GAAAT**TAATACGACTCACTATAG**GCCAAGATATAGTTGGCCAGGTTTAAGAGCTATGCTGGAAACA |
| sgHPV FWD31 | GAAAT**TAATACGACTCACTATAG**ACACAGCTATCAGAGCAAGAGTTTAAGAGCTATGCTGGAAACA |
| sgHPV FWD32 | GAAAT**TAATACGACTCACTATAG**GACTCATTCTTTGGCTAGTTTAAGAGCTATGCTGGAAACA |
| sgHPV FWD33 | GAAAT**TAATACGACTCACTATAG**GTAAAGAATAAACAATAGATGTTTAAGAGCTATGCTGGAAACA |
| sgHPV FWD34 | GAAAT**TAATACGACTCACTATAG**AGGAACAAAGGAATCGAGGGGTTTAAGAGCTATGCTGGAAACA |
| sgHPV FWD35 | GAAAT**TAATACGACTCACTATAG**GGAACAAAAGAACAAAAAAGGTTTAAGAGCTATGCTGGAAACA |

| | |
|---|---|
| sgHPV FWD36 | GAAAT**TAATACGACTCACTATAG**GTTTCCCTATGAAAATGGCAAGTTTAAGAGCTATGCTGGAAACA |

**Bold**, T7 promoter; underlined, sequence of sgRNA target; FWD, forward primer.

**Supplementary Table 3. Primers for *in vitro* T7 transcribed gRNAs targeting *MYC***

| Primer name | Sequence 5’-3’ |
|---|---|
| sgMYC FWD1 | GAAAT**TAATACGACTCACTATAG**GACCAAGTCTTGCTTACTGGTGTTTAAGAGCTATGCTGGAAACA |
| sgMYC FWD2 | GAAAT**TAATACGACTCACTATAG**GCTAAAGTACTCAAAGCAGGTTTAAGAGCTATGCTGGAAACA |
| sgMYC FWD3 | GAAAT**TAATACGACTCACTATAG**GTTTAGGCAGGGCGAGGGGGGTTTAAGAGCTATGCTGGAAACA |
| sgMYC FWD4 | GAAAT**TAATACGACTCACTATAG**GCTTGGATAACTTCTTGCAGGTTTAAGAGCTATGCTGGAAACA |
| sgMYC FWD5 | GAAAT**TAATACGACTCACTATAG**GATGCCAAAGTCAGGCTAGTTTAAGAGCTATGCTGGAAACA |
| sgMYC FWD6 | GAAAT**TAATACGACTCACTATAG**GAACAAAGAATCTGTTATAGTTTAAGAGCTATGCTGGAAACA |
| sgMYC FWD7 | GAAAT**TAATACGACTCACTATAG**GTTAATAAAAGCTGACTTCACGTTTAAGAGCTATGCTGGAAACA |
| sgMYC FWD8 | GAAAT**TAATACGACTCACTATAG**GGAACTCATTGAAGTTCGTTTAAGAGCTATGCTGGAAACA |
| sgMYC FWD9 | GAAAT**TAATACGACTCACTATAG**ATAAAACTGAAATAATTAATGTTTAAGAGCTATGCTGGAAACA |

sgMYC FWD10 GAAAT**TAATACGACTCACTATAG**ACTCACCCAAAAAACCA
GCTGTTTAAGAGCTATGCTGGAAACA

sgMYC FWD11 GAAAT**TAATACGACTCACTATAG**GAATGGTAAAGGACAGG
ATGTTTAAGAGCTATGCTGGAAACA

sgMYC FWD12 GAAAT**TAATACGACTCACTATAG**GCTTCACCAGAGAAGCA
GAGTTTAAGAGCTATGCTGGAAACA

sgMYC FWD13 GAAAT**TAATACGACTCACTATAG**GTTGATTTGGTAATAGG
CAGGTTTAAGAGCTATGCTGGAAACA

sgMYC FWD14 GAAAT**TAATACGACTCACTATAG**GACATTGGAAACTGGCC
AGTTTAAGAGCTATGCTGGAAACA

sgMYC FWD15 GAAAT**TAATACGACTCACTATAG**GCACCCTCCAGAGCTGC
AGCGGTTTAAGAGCTATGCTGGAAACA

sgMYC FWD16 GAAAT**TAATACGACTCACTATAG**GTTCTAAATACATTTGG
ACGGGTTTAAGAGCTATGCTGGAAACA

sgMYC FWD17 GAAAT**TAATACGACTCACTATAG**GTCCCAGTCTGCAAAAT
AAAGGTTTAAGAGCTATGCTGGAAACA

sgMYC FWD18 GAAAT**TAATACGACTCACTATAG**GGCTGTACCCCCAGTGA
TAGTTTAAGAGCTATGCTGGAAACA

sgMYC FWD19 GAAAT**TAATACGACTCACTATAG**ACTGTTAACCTTAACATC
AACGTTTAAGAGCTATGCTGGAAACA

sgMYC FWD20 GAAAT**TAATACGACTCACTATAG**GGTGCACAGGAGAAGAA
ATGTTTAAGAGCTATGCTGGAAACA

sgMYC FWD21 GAAAT**TAATACGACTCACTATAG**ACAAATCCTTGTCCTCCA
AGGTTTAAGAGCTATGCTGGAAACA

sgMYC FWD22 GAAAT**TAATACGACTCACTATAG**GTCCTGGGGCATAATGC
CAAGGTTTAAGAGCTATGCTGGAAACA

sgMYC FWD23 GAAAT**TAATACGACTCACTATAG**GACCAGAGCAAATTTCCTTAGTTTAAGAGCTATGCTGGAAACA

sgMYC FWD24 GAAAT**TAATACGACTCACTATAG**GCTTCCTTCTTCAATTCAGATGTTTAAGAGCTATGCTGGAAACA

sgMYC FWD25 GAAAT**TAATACGACTCACTATAG**AGAGGGATTATAAAGTTGCGGTTTAAGAGCTATGCTGGAAACA

sgMYC FWD26 GAAAT**TAATACGACTCACTATAG**ACACTGTTGTTGAAGTGGTTTAAGAGCTATGCTGGAAACA

sgMYC FWD27 GAAAT**TAATACGACTCACTATAG**ATCGTTGGAGCAAGGGTGACGGTTTAAGAGCTATGCTGGAAACA

sgMYC FWD28 GAAAT**TAATACGACTCACTATAG**GCTTCGGTTCCATCAATGTTTAAGAGCTATGCTGGAAACA

sgMYC FWD29 GAAAT**TAATACGACTCACTATAG**GTTAATCGTATATAGAGAGAAGTTTAAGAGCTATGCTGGAAACA

sgMYC FWD30 GAAAT**TAATACGACTCACTATAG**GAGAGAGCTTCTGAGCTGAGTTTAAGAGCTATGCTGGAAACA

sgMYC FWD31 GAAAT**TAATACGACTCACTATAG**ACTCTGTGAATGGCACCTTGGTTTAAGAGCTATGCTGGAAACA

sgMYC FWD32 GAAAT**TAATACGACTCACTATAG**AGAAGCTTCTGGGGTCAGTAGTTTAAGAGCTATGCTGGAAACA

sgMYC FWD33 GAAAT**TAATACGACTCACTATAG**GGGATGGTTTAAACTCTCAAGTTTAAGAGCTATGCTGGAAACA

sgMYC FWD34 GAAAT**TAATACGACTCACTATAG**GGATCATGAAAAGCTCCTACGTTTAAGAGCTATGCTGGAAACA

sgMYC FWD35 GAAAT**TAATACGACTCACTATAG**GGCAGGCTGGCAGCCTGAGTTTAAGAGCTATGCTGGAAACA sgMYC FWD36 GAAAT**TAATACGACTCACTATAG**GTTTCCCAACACACTGCTGCGTTTAAGAGCTATGCTGGAAACA

sgMYC FWD37 GAAAT**TAATACGACTCACTATAG**AAAGCGTAAATCAACAACCGTTTAAGAGCTATGCTGGAAACA

sgMYC FWD38 GAAAT**TAATACGACTCACTATAG**GGTGTGGTCCTGGAAACCCAGTTTAAGAGCTATGCTGGAAACA

sgMYC FWD39 GAAAT**TAATACGACTCACTATAG**ATATGAATTCAGTGAAAGGGTTTAAGAGCTATGCTGGAAACA

sgMYC FWD40 GAAAT**TAATACGACTCACTATAG**AAAATTCTGGAAGTTCAGGAGTTTAAGAGCTATGCTGGAAACA

sgMYC FWD41 GAAAT**TAATACGACTCACTATAG**GCAAGAGTTGGAGACAGGAAGTTTAAGAGCTATGCTGGAAACA

sgMYC FWD42 GAAAT**TAATACGACTCACTATAG**AATACAAGCTTGTCAAGTTGGTTTAAGAGCTATGCTGGAAACA

sgMYC FWD43 GAAAT**TAATACGACTCACTATAG**GGTAAGAGAATGAAGTCAATGTTTAAGAGCTATGCTGGAAACA

sgMYC FWD44 GAAAT**TAATACGACTCACTATAG**ACATTCATTCACATTAACTATGTTTAAGAGCTATGCTGGAAACA

sgMYC FWD45 GAAAT**TAATACGACTCACTATAG**GAAGAAGAGAAAAAGGGTTTAAGAGCTATGCTGGAAACA

sgMYC FWD46 GAAAT**TAATACGACTCACTATAG**GGATTCTACCCTGAGAACAAGTTTAAGAGCTATGCTGGAAACA

sgMYC FWD47 GAAAT**TAATACGACTCACTATAG**GATTCTGTAAGATGATCATGTTTAAGAGCTATGCTGGAAACA

sgMYC FWD48 GAAAT**TAATACGACTCACTATAG**GGGCTGGGAGGAAAGGGTTTAAGAGCTATGCTGGAAACA

| | |
|---|---|
| sgMYC FWD49 | GAAAT**TAATACGACTCACTATAG**GAGAAGAGACAGTAAGTCTGGTTTAAGAGCTATGCTGGAAACA |
| sgMYC FWD50 | GAAAT**TAATACGACTCACTATAG**GAAACTTGTGTGTCACTCAGAGGTTTAAGAGCTATGCTGGAAACA |
| sgMYC FWD51 | GAAAT**TAATACGACTCACTATAG**GAAATAGGGTAAGATAAGGAGGGTTTAAGAGCTATGCTGGAAACA |
| sgMYC FWD52 | GAAAT**TAATACGACTCACTATAG**ATTCTTCAGTTGACCCCCAGTTTAAGAGCTATGCTGGAAACA |
| sgMYC FWD53 | GAAAT**TAATACGACTCACTATAG**GGAAAATTAATCAGTGTGCAGTTTAAGAGCTATGCTGGAAACA |
| sgMYC FWD54 | GAAAT**TAATACGACTCACTATAG**GTCCATGTAGAAGACGTTACAGGTTTAAGAGCTATGCTGGAAACA |
| sgMYC FWD55 | GAAAT**TAATACGACTCACTATAG**ATAAAAGTGAAAACAACCAAGTTTAAGAGCTATGCTGGAAACA |
| sgMYC FWD56 | GAAAT**TAATACGACTCACTATAG**GCAGAGATCCCAAGCTAGTTTAAGAGCTATGCTGGAAACA |
| sgMYC FWD57 | GAAAT**TAATACGACTCACTATAG**AAGCAAAAGTTTCTCTCCAAGTTTAAGAGCTATGCTGGAAACA |
| sgMYC FWD58 | GAAAT**TAATACGACTCACTATAG**GCTGTATGTCCAACCACGCAAGTTTAAGAGCTATGCTGGAAACA |
| sgMYC FWD59 | GAAAT**TAATACGACTCACTATAG**GGCCAGTGAGCCAAAGTGGTTTAAGAGCTATGCTGGAAACA |
| sgMYC FWD60 | GAAAT**TAATACGACTCACTATAG**GTTGAACCCCTTCAAAGGCTGTTTAAGAGCTATGCTGGAAACA |
| sgMYC FWD61 | GAAAT**TAATACGACTCACTATAG**AGTAGCTACAAAGTAAGGCAGTTTAAGAGCTATGCTGGAAACA |

| | |
|---|---|
| sgMYC FWD62 | GAAAT**TAATACGACTCACTATAG**ATAAAAGAAGGCCAAGCTGGGTTTAAGAGCTATGCTGGAAACA |
| sgMYC FWD63 | GAAAT**TAATACGACTCACTATAG**GTCCATTCTGAGACCTTCTGCTGTTTAAGAGCTATGCTGGAAACA |
| sgMYC FWD64 | GAAAT**TAATACGACTCACTATAG**GGATATGGAAGCTCTCGTTTAAGAGCTATGCTGGAAACA |
| sgMYC FWD65 | GAAAT**TAATACGACTCACTATAG**GCAGTATACTAAAAGCCAGGGTTTAAGAGCTATGCTGGAAACA |
| sgMYC FWD66 | GAAAT**TAATACGACTCACTATAG**GCATAGGAATGATAACAAAAAGTTTAAGAGCTATGCTGGAAACA |
| sgMYC FWD67 | GAAAT**TAATACGACTCACTATAG**GTTCCAACGAGAGGAACACGTTTAAGAGCTATGCTGGAAACA |
| sgMYC FWD68 | GAAAT**TAATACGACTCACTATAG**GATTAAAAACCAAGCTAGCCAGTTTAAGAGCTATGCTGGAAACA |
| sgMYC FWD69 | GAAAT**TAATACGACTCACTATAG**GATAAATATCTATCTCCAGGTTTAAGAGCTATGCTGGAAACA |
| sgMYC FWD70 | GAAAT**TAATACGACTCACTATAG**GACAGGCCGCACGTGACTTGAGTTTAAGAGCTATGCTGGAAACA |
| sgMYC FWD71 | GAAAT**TAATACGACTCACTATAG**GGTGAGACTGAAGCCCCTCAGTTTAAGAGCTATGCTGGAAACA |
| sgMYC FWD72 | GAAAT**TAATACGACTCACTATAG**GATGGAAAGCAACTGGACGGTTTAAGAGCTATGCTGGAAACA |
| sgMYC FWD73 | GAAAT**TAATACGACTCACTATAG**GATGTTCAAAGAAGGTGTTGGTTTAAGAGCTATGCTGGAAACA |
| sgMYC FWD74 | GAAAT**TAATACGACTCACTATAG**GTTCTCCCCAGAGACACAAAAGTTTAAGAGCTATGCTGGAAACA |

sgMYC FWD75 GAAAT**TAATACGACTCACTATAG**GTCATATATTTTTATACTTGTTTAAGAGCTATGCTGGAAACA

sgMYC FWD76 GAAAT**TAATACGACTCACTATAG**ACACACTATTCTGTTCTGTGGTTTAAGAGCTATGCTGGAAACA

sgMYC FWD77 GAAAT**TAATACGACTCACTATAG**GCAGGTCTCCTGGAGGGCCGGTTTAAGAGCTATGCTGGAAACA

sgMYC FWD78 GAAAT**TAATACGACTCACTATAG**GCAGCAGCTCCAAATAACAGGTTTAAGAGCTATGCTGGAAACA

sgMYC FWD79 GAAAT**TAATACGACTCACTATAG**GCTGCTAGAGCAACAAGCAAGTTTAAGAGCTATGCTGGAAACA

sgMYC FWD80 GAAAT**TAATACGACTCACTATAG**GGTCTTCCAAAAAAAATTGGTTTAAGAGCTATGCTGGAAACA

sgMYC FWD81 GAAAT**TAATACGACTCACTATAG**AATACTGACCAGTCAGGTTTAAGAGCTATGCTGGAAACA

sgMYC FWD82 GAAAT**TAATACGACTCACTATAG**GCCAATGGACACGTATCACTTGTTTAAGAGCTATGCTGGAAACA

sgMYC FWD83 GAAAT**TAATACGACTCACTATAG**GACATTTGCTGGGTTGAAAAAGTTTAAGAGCTATGCTGGAAACA

sgMYC FWD84 GAAAT**TAATACGACTCACTATAG**ATGTCCTTTAACCTGGCTGGTTTAAGAGCTATGCTGGAAACA

sgMYC FWD85 GAAAT**TAATACGACTCACTATAG**AGGTGATGTCACCAGCCTGAGTTTAAGAGCTATGCTGGAAACA

sgMYC FWD86 GAAAT**TAATACGACTCACTATAG**ACAGGCATTATATCTGCCTGGTTTAAGAGCTATGCTGGAAACA

sgMYC FWD87 GAAAT**TAATACGACTCACTATAG**GGCGGATATACCACATCTTGTTTAAGAGCTATGCTGGAAACA

| | |
|---|---|
| sgMYC FWD88 | GAAAT**TAATACGACTCACTATAG**ACAATGGCAAAACCAAAAGTGTTTAAGAGCTATGCTGGAAACA |
| sgMYC FWD89 | GAAAT**TAATACGACTCACTATAG**GCTAAATGAGTGCTCTCCACAGTTTAAGAGCTATGCTGGAAACA |
| sgMYC FWD90 | GAAAT**TAATACGACTCACTATAG**GTCAGCCTACAAGGCTCCTGCGTTTAAGAGCTATGCTGGAAACA |
| sgMYC FWD91 | GAAAT**TAATACGACTCACTATAG**GTCCACTGCCAGAAGTCCTTAGTTTAAGAGCTATGCTGGAAACA |
| sgMYC FWD92 | GAAAT**TAATACGACTCACTATAG**GTAAAAACCTACTTGACCAGTTTAAGAGCTATGCTGGAAACA |
| sgMYC FWD93 | GAAAT**TAATACGACTCACTATAG**ACTAAAATGAGTATGCAATAGTTTAAGAGCTATGCTGGAAACA |
| sgMYC FWD94 | GAAAT**TAATACGACTCACTATAG**GATCTCATAGAAAAAAAGTGTGAGTTTAAGAGCTATGCTGGAAACA |
| sgMYC FWD95 | GAAAT**TAATACGACTCACTATAG**GATGAGTTTCTAAGACGTGGGTTTAAGAGCTATGCTGGAAACA |
| sgMYC FWD96 | GAAAT**TAATACGACTCACTATAG**AATTCCAACAAACCCTAAAAGTTTAAGAGCTATGCTGGAAACA |
| sgMYC FWD97 | GAAAT**TAATACGACTCACTATAG**GCCATTACCGGTTCTCCATAGTTTAAGAGCTATGCTGGAAACA |
| sgMYC FWD98 | GAAAT**TAATACGACTCACTATAG**GGAGTTACTGGAGGAAAAAGGTTTAAGAGCTATGCTGGAAACA |
| sgMYC FWD99 | GAAAT**TAATACGACTCACTATAG**AACCTGAAAGAATAACAAGGGTTTAAGAGCTATGCTGGAAACA |
| sgMYC FWD100 | GAAAT**TAATACGACTCACTATAG**GCTGGAAACCTTGCACCTGTTTAAGAGCTATGCTGGAAACA |

sgMYC FWD101 GAAAT**TAATACGACTCACTATAG**ATCGCGCCTGGATGTCAACGAGTTTAAGAGCTATGCTGGAAACA

sgMYC FWD102 GAAAT**TAATACGACTCACTATAG**GTACTTTCGCAAACCTGAACGGTTTAAGAGCTATGCTGGAAACA

sgMYC FWD103 GAAAT**TAATACGACTCACTATAG**AGGCCTTTGCCGCAAACGCGGTTTAAGAGCTATGCTGGAAACA

sgMYC FWD104 GAAAT**TAATACGACTCACTATAG**GCTGAATTGTGCAGTGCATGTTTAAGAGCTATGCTGGAAACA

sgMYC FWD105 GAAAT**TAATACGACTCACTATAG**GAACGCTGAGCTGCAAACTCAAGTTTAAGAGCTATGCTGGAAACA

sgMYC FWD106 GAAAT**TAATACGACTCACTATAG**GCATGTACGCTGTTCAAGATGTTTAAGAGCTATGCTGGAAACA

sgMYC FWD107 GAAAT**TAATACGACTCACTATAG**GCAAAAGAGAAAACAATTCGGGTTTAAGAGCTATGCTGGAAACA

sgMYC FWD108 GAAAT**TAATACGACTCACTATAG**ATCCTTGGTCCCTCACCCAAGTTTAAGAGCTATGCTGGAAACA

sgMYC FWD109 GAAAT**TAATACGACTCACTATAG**GCACAAAATAAAAAATCCCGAGTTTAAGAGCTATGCTGGAAACA

sgMYC FWD110 GAAAT**TAATACGACTCACTATAG**GAGCAAACAAATCATGTGTGGTTTAAGAGCTATGCTGGAAACA

sgMYC FWD111 GAAAT**TAATACGACTCACTATAG**GTGAATACACGTTTGCGTTTAAGAGCTATGCTGGAAACA

sgMYC FWD112 GAAAT**TAATACGACTCACTATAG**GTGAACTAGGAAATTAATGCCGTTTAAGAGCTATGCTGGAAACA

sgMYC FWD113 GAAAT**TAATACGACTCACTATAG**GCCCCCCCCCAAAAAAAGGCAGTTTAAGAGCTATGCTGGAAACA sgMYC FWD114 GAAAT**TAATACGACTCACTATAG**ATCGATTCTGATCAAAGAAGGTTTAAGAGCTATGCTGGAAACA

sgMYC FWD115 GAAAT**TAATACGACTCACTATAG**GACCGCATTTCCAATAATAAAAGTTTAAGAGCTATGCTGGAAACA

sgMYC FWD116 GAAAT**TAATACGACTCACTATAG**GTTAAACGTCCGGTTTGTCCGGTTTAAGAGCTATGCTGGAAACA

sgMYC FWD117 GAAAT**TAATACGACTCACTATAG**AGAGCTTGTGGACCGAGCCGGTTTAAGAGCTATGCTGGAAACA

sgMYC FWD118 GAAAT**TAATACGACTCACTATAG**GTAGACGGGAGAATATGGGAGGTTTAAGAGCTATGCTGGAAACA

sgMYC FWD119 GAAAT**TAATACGACTCACTATAG**GTTGCAAACCGGCGCCACAGTTTAAGAGCTATGCTGGAAACA

sgMYC FWD120 GAAAT**TAATACGACTCACTATAG**GAGAAATTGGGAACTCCGTGGTTTAAGAGCTATGCTGGAAACA

sgMYC FWD121 GAAAT**TAATACGACTCACTATAG**GCTGGGCTAGGGCGAGAGGGGTTTAAGAGCTATGCTGGAAACA

sgMYC FWD122 GAAAT**TAATACGACTCACTATAG**GCTAAACAGACGCCTCCCGCAGTTTAAGAGCTATGCTGGAAACA

sgMYC FWD123 GAAAT**TAATACGACTCACTATAG**GGGGGACTCAGTCTGGGGTTTAAGAGCTATGCTGGAAACA

sgMYC FWD124 GAAAT**TAATACGACTCACTATAG**GACTCCCCCCAACAAATGCAAGTTTAAGAGCTATGCTGGAAACA

sgMYC FWD125 GAAAT**TAATACGACTCACTATAG**ACGCGCTCTCCAAGTATACGGTTTAAGAGCTATGCTGGAAACA

sgMYC FWD126 GAAAT**TAATACGACTCACTATAG**GGAATGATAGAGGCATAAGGGTTTAAGAGCTATGCTGGAAACA

| | |
|---|---|
| sgMYC FWD127 | GAAAT**TAATACGACTCACTATAG**GGGCGCGCGTTCAGAGCGTGTTTAAGAGCTATGCTGGAAACA |
| sgMYC FWD128 | GAAAT**TAATACGACTCACTATAG**GGGACTCTTGATCAAAGCGGTTTAAGAGCTATGCTGGAAACA |
| sgMYC FWD129 | GAAAT**TAATACGACTCACTATAG**GCAGCCTGGTACGCGCGGTTTAAGAGCTATGCTGGAAACA |
| sgMYC FWD130 | GAAAT**TAATACGACTCACTATAG**ATACTCACAGGACAAGGATGGTTTAAGAGCTATGCTGGAAACA |
| sgMYC FWD131 | GAAAT**TAATACGACTCACTATAG**GGAGCAGCAGAGAAAGGGAGGTTTAAGAGCTATGCTGGAAACA |
| sgMYC FWD132 | GAAAT**TAATACGACTCACTATAG**GCGCGCGTAGTTAATTCATGGTTTAAGAGCTATGCTGGAAACA |
| sgMYC FWD133 | GAAAT**TAATACGACTCACTATAG**GGTGGGGAGGAGACTCAGCCGTTTAAGAGCTATGCTGGAAACA |
| sgMYC FWD134 | GAAAT**TAATACGACTCACTATAG**GGGTTCCCAAAGCAGAGTTTAAGAGCTATGCTGGAAACA |
| sgMYC FWD135 | GAAAT**TAATACGACTCACTATAG**GTTATAATGCGAGGGTCTGGAGTTTAAGAGCTATGCTGGAAACA |
| sgMYC FWD136 | GAAAT**TAATACGACTCACTATAG**GAGAAGGGCAGGGCTTCTCAGGTTTAAGAGCTATGCTGGAAACA |
| sgMYC FWD137 | GAAAT**TAATACGACTCACTATAG**GGGAAAAAGAACGGAGGGAGTTTAAGAGCTATGCTGGAAACA |
| sgMYC FWD138 | GAAAT**TAATACGACTCACTATAG**GCTGTAGTAATTCCAGCGAGGTTTAAGAGCTATGCTGGAAACA |
| sgMYC FWD139 | GAAAT**TAATACGACTCACTATAG**GGGCGAGCAGAGCTGCGCTGGTTTAAGAGCTATGCTGGAAACA |

| | |
|---|---|
| sgMYC FWD140 | GAAAT**TAATACGACTCACTATAG**GGAGATCCGGAGCGAATAGGGTTTAAGAGCTATGCTGGAAACA |
| sgMYC FWD141 | GAAAT**TAATACGACTCACTATAG**ACCGCTGGCTGGGGATCAGGTTTAAGAGCTATGCTGGAAACA |
| sgMYC FWD142 | GAAAT**TAATACGACTCACTATAG**GAAACTTTGCCCATAGCAGGTTTAAGAGCTATGCTGGAAACA |
| sgMYC FWD143 | GAAAT**TAATACGACTCACTATAG**GACGCGACTCTCCCGACGCGGTTTAAGAGCTATGCTGGAAACA |
| sgMYC FWD144 | GAAAT**TAATACGACTCACTATAG**GCGGGTCCTGGCAGCGGCGGTTTAAGAGCTATGCTGGAAACA |
| sgMYC FWD145 | GAAAT**TAATACGACTCACTATAG**GCAGCTGCTTAGACGCGTTTAAGAGCTATGCTGGAAACA |
| sgMYC FWD146 | GAAAT**TAATACGACTCACTATAG**GATGAGTCGAATGCCTAAATAGTTTAAGAGCTATGCTGGAAACA |
| sgMYC FWD147 | GAAAT**TAATACGACTCACTATAG**GAGAAAAGTGTCAATAGCGCGTTTAAGAGCTATGCTGGAAACA |
| sgMYC FWD148 | GAAAT**TAATACGACTCACTATAG**GGTAATCCAGAACTGGATCGGTTTAAGAGCTATGCTGGAAACA |
| sgMYC FWD149 | GAAAT**TAATACGACTCACTATAG**GATGGGAGAGGAGAAGGCAGGTTTAAGAGCTATGCTGGAAACA |
| sgMYC FWD150 | GAAAT**TAATACGACTCACTATAG**ATAAGGCAGAAATCTCGAAAGTTTAAGAGCTATGCTGGAAACA |
| sgMYC FWD151 | GAAAT**TAATACGACTCACTATAG**ATAAAGCAGGAATGTCCGACGTTTAAGAGCTATGCTGGAAACA |
| sgMYC FWD152 | GAAAT**TAATACGACTCACTATAG**GCTGGGGGTTGCTTTGCGGTGTTTAAGAGCTATGCTGGAAACA |

| | |
|---|---|
| sgMYC FWD153 | GAAAT**TAATACGACTCACTATAG**GGCTCACACAGGCGATATGGTTTAAGAGCTATGCTGGAAACA |
| sgMYC FWD154 | GAAAT**TAATACGACTCACTATAG**GACTTGTCCCCGTCTCCGGGGTTTAAGAGCTATGCTGGAAACA |
| sgMYC FWD155 | GAAAT**TAATACGACTCACTATAG**ACAGCCGGAGACGGACACTGGTTTAAGAGCTATGCTGGAAACA |
| sgMYC FWD156 | GAAAT**TAATACGACTCACTATAG**GGCGGGTTGGAATCGCCGCGGTTTAAGAGCTATGCTGGAAACA |
| sgMYC FWD157 | GAAAT**TAATACGACTCACTATAG**ATTTAAAACCTGGGTCTCTAGGTTTAAGAGCTATGCTGGAAACA |
| sgMYC FWD158 | GAAAT**TAATACGACTCACTATAG**GTGTTGGGTAGGCGCAGGCAGGTTTAAGAGCTATGCTGGAAACA |
| sgMYC FWD159 | GAAAT**TAATACGACTCACTATAG**GTCGTTGACTTGGAAAAACCAGTTTAAGAGCTATGCTGGAAACA |
| sgMYC FWD160 | GAAAT**TAATACGACTCACTATAG**AGCCCTGACTCCCCTGCCGGTTTAAGAGCTATGCTGGAAACA |
| sgMYC FWD161 | GAAAT**TAATACGACTCACTATAG**GAGATGCGGAGGAACTGCGGTTTAAGAGCTATGCTGGAAACA |
| sgMYC FWD162 | GAAAT**TAATACGACTCACTATAG**GCCCCGGAGCACCCCACCAAGTTTAAGAGCTATGCTGGAAACA |
| sgMYC FWD163 | GAAAT**TAATACGACTCACTATAG**GCATCTCCGTATTGAGTGCGAAGTTTAAGAGCTATGCTGGAAACA |
| sgMYC FWD164 | GAAAT**TAATACGACTCACTATAG**GGAGGGGTGTTAAAGCCCGGTTTAAGAGCTATGCTGGAAACA |
| sgMYC FWD165 | GAAAT**TAATACGACTCACTATAG**GGAGAAGGCGAGAGGCGCCTGTTTAAGAGCTATGCTGGAAACA |

sgMYC FWD166 GAAAT**TAATACGACTCACTATAG**GAAAACAATTTGCCAAA
ATCCAGTTTAAGAGCTATGCTGGAAACA

sgMYC FWD167 GAAAT**TAATACGACTCACTATAG**GCGGCTTCTTAAGGGCG
CCAGTTTAAGAGCTATGCTGGAAACA

sgMYC FWD168 GAAAT**TAATACGACTCACTATAG**GCGCTCCGGGCTCCCGG
GTTTAAGAGCTATGCTGGAAACA

sgMYC FWD169 GAAAT**TAATACGACTCACTATAG**GTGCGTCTCCGAGATAG
CAGGTTTAAGAGCTATGCTGGAAACA

sgMYC FWD170 GAAAT**TAATACGACTCACTATAG**GGTCTTGGTGGGGGAAT
AAAGTTTAAGAGCTATGCTGGAAACA

sgMYC FWD171 GAAAT**TAATACGACTCACTATAG**GGGGAGAGGTTCGGGAC
TGGTTTAAGAGCTATGCTGGAAACA

sgMYC FWD172 GAAAT**TAATACGACTCACTATAG**GAGGCAGTCTTGAGTTA
AAGGTTTAAGAGCTATGCTGGAAACA

sgMYC FWD173 GAAAT**TAATACGACTCACTATAG**GTTGGTGAAGCTAACGT
TGAGTTTAAGAGCTATGCTGGAAACA

sgMYC FWD174 GAAAT**TAATACGACTCACTATAG**GTATTTCTACTGCGACG
AGGGTTTAAGAGCTATGCTGGAAACA

sgMYC FWD175 GAAAT**TAATACGACTCACTATAG**GATATCCTCGCTGGGCG
CCGGTTTAAGAGCTATGCTGGAAACA

sgMYC FWD176 GAAAT**TAATACGACTCACTATAG**GCGAGCAGAGCCCGGAG
CGGGTTTAAGAGCTATGCTGGAAACA

sgMYC FWD177 GAAAT**TAATACGACTCACTATAG**GCTTCGGGGAGACAACG
ACGGGTTTAAGAGCTATGCTGGAAACA

sgMYC FWD178 GAAAT**TAATACGACTCACTATAG**GCTCGGTCACCATCTCC
AGCGTTTAAGAGCTATGCTGGAAACA

sgMYC FWD179 GAAAT**TAATACGACTCACTATAG**ATCATCATCCAGGACTGTATGGTTTAAGAGCTATGCTGGAAACA

sgMYC FWD180 GAAAT**TAATACGACTCACTATAG**GCTCTGAGACGAGCTTGGCGGGTTTAAGAGCTATGCTGGAAACA

sgMYC FWD181 GAAAT**TAATACGACTCACTATAG**GGCTGCGCGCAAAGACAGGTTTAAGAGCTATGCTGGAAACA

sgMYC FWD182 GAAAT**TAATACGACTCACTATAG**GGAGCAGACGCTGTGGCCGGTTTAAGAGCTATGCTGGAAACA

sgMYC FWD183 GAAAT**TAATACGACTCACTATAG**GGTAGGGGAAGACCACCGAGGTTTAAGAGCTATGCTGGAAACA

sgMYC FWD184 GAAAT**TAATACGACTCACTATAG**GCTGGAGTCTTGCGAGGCGCGTTTAAGAGCTATGCTGGAAACA

sgMYC FWD185 GAAAT**TAATACGACTCACTATAG**GAGGAGAGCAGAGAATCCGGTTTAAGAGCTATGCTGGAAACA

sgMYC FWD186 GAAAT**TAATACGACTCACTATAG**GTCTCCTCATGGAGCACCAGTTTAAGAGCTATGCTGGAAACA

sgMYC FWD187 GAAAT**TAATACGACTCACTATAG**GGCCTGTCAAAAGTGGGGTTTAAGAGCTATGCTGGAAACA

sgMYC FWD188 GAAAT**TAATACGACTCACTATAG**AATAAGCTGCCAATGAAAATGTTTAAGAGCTATGCTGGAAACA

sgMYC FWD189 GAAAT**TAATACGACTCACTATAG**GCTAAAGCCCAAGGTTTCAGGTTTAAGAGCTATGCTGGAAACA

sgMYC FWD190 GAAAT**TAATACGACTCACTATAG**GCAAACATGGGCAGTCTAAGGTTTAAGAGCTATGCTGGAAACA

sgMYC FWD191 GAAAT**TAATACGACTCACTATAG**GAGTTGTAAGATAAGCCAGAGTTTAAGAGCTATGCTGGAAACA sgMYC FWD192 GAAAT**TAATACGACTCACTATAG**GCAATTAAAATGTTAACGGGGTTTAAGAGCTATGCTGGAAACA

sgMYC FWD193 GAAAT**TAATACGACTCACTATAG**GTATGAATGAGGATAAGAGGTTTAAGAGCTATGCTGGAAACA

sgMYC FWD194 GAAAT**TAATACGACTCACTATAG**GCCACTTCTCGGAAGTTAAGAGTTTAAGAGCTATGCTGGAAACA

sgMYC FWD195 GAAAT**TAATACGACTCACTATAG**GTGTTTAGAGGCTAGGCAGTTTAAGAGCTATGCTGGAAACA

sgMYC FWD196 GAAAT**TAATACGACTCACTATAG**GAACTGCCTCAAGAGTGGGTGTTTAAGAGCTATGCTGGAAACA

sgMYC FWD197 GAAAT**TAATACGACTCACTATAG**GCCAAAAGTCCAAGAGGGCGGGTTTAAGAGCTATGCTGGAAACA

sgMYC FWD198 GAAAT**TAATACGACTCACTATAG**AATGATAGCTGCAAATTGCTGTTTAAGAGCTATGCTGGAAACA

sgMYC FWD199 GAAAT**TAATACGACTCACTATAG**GTAAAGTCCCTCAAAAATAGGGTTTAAGAGCTATGCTGGAAACA

sgMYC FWD200 GAAAT**TAATACGACTCACTATAG**GTCCAAAGCCTCATTAAGTCTTGTTTAAGAGCTATGCTGGAAACA

sgMYC FWD201 GAAAT**TAATACGACTCACTATAG**GACAGCTGGGTTATGGCAGTTTAAGAGCTATGCTGGAAACA

sgMYC FWD202 GAAAT**TAATACGACTCACTATAG**GCTTCATGGTGAGAGGAGTAAGTTTAAGAGCTATGCTGGAAACA

sgMYC FWD203 GAAAT**TAATACGACTCACTATAG**GTATTTGTACAGCATTAATCGTTTAAGAGCTATGCTGGAAACA

sgMYC FWD204 GAAAT**TAATACGACTCACTATAG**GAAATCACTCCTTTAGCAGTTTAAGAGCTATGCTGGAAACA sgMYC FWD205 GAAAT**TAATACGACTCACTATAG**GAGGAGGAACAAGAAGATGGTTTAAGAGCTATGCTGGAAACA

sgMYC FWD206 GAAAT**TAATACGACTCACTATAG**GTTTCTGTGGAAAAGAGGCGTTTAAGAGCTATGCTGGAAACA

sgMYC FWD207 GAAAT**TAATACGACTCACTATAG**GGCACCTCTTGAGGACCAGTGTTTAAGAGCTATGCTGGAAACA

sgMYC FWD208 GAAAT**TAATACGACTCACTATAG**GCAGGATAGTCCTTCCGAGGTTTAAGAGCTATGCTGGAAACA

sgMYC FWD209 GAAAT**TAATACGACTCACTATAG**GGTTGTTGCTGATCTGTCTCGTTTAAGAGCTATGCTGGAAACA

sgMYC FWD210 GAAAT**TAATACGACTCACTATAG**GCCTCTTGACATTCTCCTGTTTAAGAGCTATGCTGGAAACA

sgMYC FWD211 GAAAT**TAATACGACTCACTATAG**GAGGAGGAACGAGCTAAAAGTTTAAGAGCTATGCTGGAAACA

sgMYC FWD212 GAAAT**TAATACGACTCACTATAG**GGAGTTGGAAAACAATGAAAGTTTAAGAGCTATGCTGGAAACA

sgMYC FWD213 GAAAT**TAATACGACTCACTATAG**ACATCCTGTCCGTCCAAGCAGGTTTAAGAGCTATGCTGGAAACA

sgMYC FWD214 GAAAT**TAATACGACTCACTATAG**GTGCGTAAGGAAAAGTAGTTTAAGAGCTATGCTGGAAACA

sgMYC FWD215 GAAAT**TAATACGACTCACTATAG**GCATTTGAAACAAGTTCATGTTTAAGAGCTATGCTGGAAACA

sgMYC FWD216 GAAAT**TAATACGACTCACTATAG**GTCTCAAGACTCAGCCAGTTTAAGAGCTATGCTGGAAACA

sgMYC FWD217 GAAAT**TAATACGACTCACTATAG**ACATTCACAACTTAAGATTGTTTAAGAGCTATGCTGGAAACA sgMYC FWD218 GAAAT**TAATACGACTCACTATAG**GCAATTGATGAAAACAAACAGTTTAAGAGCTATGCTGGAAACA

sgMYC FWD219 GAAAT**TAATACGACTCACTATAG**GAGTTTTCCTCTGTTGAAATGTTTAAGAGCTATGCTGGAAACA

sgMYC FWD220 GAAAT**TAATACGACTCACTATAG**GAGGTTCTAAGATGCTTCCGTTTAAGAGCTATGCTGGAAACA

sgMYC FWD221 GAAAT**TAATACGACTCACTATAG**GTAGGCAAAGGAGATACAAGGTTTAAGAGCTATGCTGGAAACA

sgMYC FWD222 GAAAT**TAATACGACTCACTATAG**GGGAGTTGGGAGGAAGGTGGTTTAAGAGCTATGCTGGAAACA

sgMYC FWD223 GAAAT**TAATACGACTCACTATAG**ATTCCTGGGTTTGGAGTGAGCAGTTTAAGAGCTATGCTGGAAACA

sgMYC FWD224 GAAAT**TAATACGACTCACTATAG**ACCAAGGCATGATAGCGAAGTTTAAGAGCTATGCTGGAAACA

sgMYC FWD225 GAAAT**TAATACGACTCACTATAG**AGGTTTAGGACCTAAGTTTAAGAGCTATGCTGGAAACA

sgMYC FWD226 GAAAT**TAATACGACTCACTATAG**GTTTCCCCTTGACCATGAGGAGTTTAAGAGCTATGCTGGAAACA

sgMYC FWD227 GAAAT**TAATACGACTCACTATAG**GCTGGCTGCTTGTGAGTACGTTTAAGAGCTATGCTGGAAACA

sgMYC FWD228 GAAAT**TAATACGACTCACTATAG**AACATCTAAGCCTGGTCGTTTAAGAGCTATGCTGGAAACA

sgMYC FWD229 GAAAT**TAATACGACTCACTATAG**GGCTAAGGTAGGAGTCAAGAGTTTAAGAGCTATGCTGGAAACA

sgMYC FWD230 GAAAT**TAATACGACTCACTATAG**ATCTGAACTGGCTTCTTCCCGTTTAAGAGCTATGCTGGAAACA sgMYC FWD231 GAAAT**TAATACGACTCACTATAG**GTAAAAAAGGATGGAAGCAGTTTAAGAGCTATGCTGGAAACA

sgMYC FWD232 GAAAT**TAATACGACTCACTATAG**GTTCAAAAATACCTTTTCAGTTTAAGAGCTATGCTGGAAACA

sgMYC FWD233 GAAAT**TAATACGACTCACTATAG**ATTTCTGGTTTGGGCCATGTTTAAGAGCTATGCTGGAAACA

sgMYC FWD234 GAAAT**TAATACGACTCACTATAG**ATTGTCTCAGTCTCAAAGTGTTTAAGAGCTATGCTGGAAACA

sgMYC FWD235 GAAAT**TAATACGACTCACTATAG**ATTTATTGATTTATGGGTGGGTTTAAGAGCTATGCTGGAAACA

sgMYC FWD236 GAAAT**TAATACGACTCACTATAG**GGATCAAGAAAAAGACATTAGTTTAAGAGCTATGCTGGAAACA

sgMYC FWD237 GAAAT**TAATACGACTCACTATAG**GTCATATAGGCGAATTTCAAAGTTTAAGAGCTATGCTGGAAACA

sgMYC FWD238 GAAAT**TAATACGACTCACTATAG**GCTCAGTCTTTGCCCCTTTGGTTTAAGAGCTATGCTGGAAACA

sgMYC FWD239 GAAAT**TAATACGACTCACTATAG**GACTCACTTGGGAATCGGGAGTTTAAGAGCTATGCTGGAAACA

sgMYC FWD240 GAAAT**TAATACGACTCACTATAG**GAACACTCTCTCCTATTCTGGTTTAAGAGCTATGCTGGAAACA

sgMYC FWD241 GAAAT**TAATACGACTCACTATAG**GGAAAGAACTTTAGGGATGGTTTAAGAGCTATGCTGGAAACA

sgMYC FWD242 GAAAT**TAATACGACTCACTATAG**GGTAAGAGCGGCCTAATGTTTAAGAGCTATGCTGGAAACA

sgMYC FWD243 GAAAT**TAATACGACTCACTATAG**GCCTGACTTTCGGGAAGGAAGTGTTTAAGAGCTATGCTGGAAACA

| | |
|---|---|
| sgMYC FWD244 | GAAAT**TAATACGACTCACTATAG**GCCTATACAGGGAGTCCCAGGTTTAAGAGCTATGCTGGAAACA |
| sgMYC FWD245 | GAAAT**TAATACGACTCACTATAG**ATCAAGAATCGGACGTGAAGTTTAAGAGCTATGCTGGAAACA |
| sgMYC FWD246 | GAAAT**TAATACGACTCACTATAG**GGATAGTGTCATGGATAAAGTTTAAGAGCTATGCTGGAAACA |
| sgMYC FWD247 | GAAAT**TAATACGACTCACTATAG**GGAATGATTTTGTTGAGGGAGTTTAAGAGCTATGCTGGAAACA |
| sgMYC FWD248 | GAAAT**TAATACGACTCACTATAG**GATCTCCTTTGTTGCTTCCAAAGTTTAAGAGCTATGCTGGAAACA |
| sgMYC FWD249 | GAAAT**TAATACGACTCACTATAG**GCTCTCTAAGTATTAGGCTGTTTAAGAGCTATGCTGGAAACA |
| sgMYC FWD250 | GAAAT**TAATACGACTCACTATAG**GACAAACTCTCACACAAAACTGTTTAAGAGCTATGCTGGAAACA |
| sgMYC FWD251 | GAAAT**TAATACGACTCACTATAG**GTTTGCAATAACTATAATGTTTAAGAGCTATGCTGGAAACA |
| sgMYC FWD252 | GAAAT**TAATACGACTCACTATAG**GATGGGAAAAAATGCTACAGGTTTAAGAGCTATGCTGGAAACA |
| sgMYC FWD253 | GAAAT**TAATACGACTCACTATAG**GTGCATTTATAGACAAGGGGTTTAAGAGCTATGCTGGAAACA |
| sgMYC FWD254 | GAAAT**TAATACGACTCACTATAG**GAATATTATAAGACTACATTAAGTTTAAGAGCTATGCTGGAAACA |
| sgMYC FWD255 | GAAAT**TAATACGACTCACTATAG**GCCAGTAGGATGGGAGCAGTTTAAGAGCTATGCTGGAAACA |
| sgMYC FWD256 | GAAAT**TAATACGACTCACTATAG**GACTTTTTGCTAAGGCTTTGGGTTTAAGAGCTATGCTGGAAACA |

sgMYC FWD257 GAAAT**TAATACGACTCACTATAG**GTTTTCCTCTAAATGGAGAGTGTTTAAGAGCTATGCTGGAAACA

sgMYC FWD258 GAAAT**TAATACGACTCACTATAG**GCTTTTTCGGAAGACAGAGTTGAGTTTAAGAGCTATGCTGGAAACA

sgMYC FWD259 GAAAT**TAATACGACTCACTATAG**GGGACTGTGGCTGAGGTCCCGTTTAAGAGCTATGCTGGAAACA

sgMYC FWD260 GAAAT**TAATACGACTCACTATAG**ATAGACCTCAGATTGCACGTTTAAGAGCTATGCTGGAAACA

sgMYC FWD261 GAAAT**TAATACGACTCACTATAG**GAGTGTATGTATGTAATAAGTTTAAGAGCTATGCTGGAAACA

sgMYC FWD262 GAAAT**TAATACGACTCACTATAG**GGAGGAGGTTCAAGCTCCGTTTAAGAGCTATGCTGGAAACA

sgMYC FWD263 GAAAT**TAATACGACTCACTATAG**GAATCCAGGAAAGAGCCCGTTTAAGAGCTATGCTGGAAACA

sgMYC FWD264 GAAAT**TAATACGACTCACTATAG**GAGAGATAAGGAGAAGCTGTTTAAGAGCTATGCTGGAAACA

sgMYC FWD265 GAAAT**TAATACGACTCACTATAG**GGAAGCCCTGGTGTGTCAAGTTTAAGAGCTATGCTGGAAACA

sgMYC FWD266 GAAAT**TAATACGACTCACTATAG**ATCACATGAACGCAGCTTCCGTTTAAGAGCTATGCTGGAAACA

sgMYC FWD267 GAAAT**TAATACGACTCACTATAG**GAATGAAGGAGAGGATGCCGGGTTTAAGAGCTATGCTGGAAACA

sgMYC FWD268 GAAAT**TAATACGACTCACTATAG**GAGAAGGTAGGGCAGGGTGCTGTTTAAGAGCTATGCTGGAAACA

sgMYC FWD269 GAAAT**TAATACGACTCACTATAG**GAAGAGGATCACTGGGAATGAGTTTAAGAGCTATGCTGGAAACA sgMYC FWD270 GAAAT**TAATACGACTCACTATAG**GGAGGCAGCATAGGACTGAGGTTTAAGAGCTATGCTGGAAACA

sgMYC FWD271 GAAAT**TAATACGACTCACTATAG**AATGAAGTAGGTAGACAAAAGTTTAAGAGCTATGCTGGAAACA

sgMYC FWD272 GAAAT**TAATACGACTCACTATAG**GCAGGAATGATTCAATCTAGGTTTAAGAGCTATGCTGGAAACA

sgMYC FWD273 GAAAT**TAATACGACTCACTATAG**GTGATGGTGCTACCAACCGAAGTTTAAGAGCTATGCTGGAAACA

sgMYC FWD274 GAAAT**TAATACGACTCACTATAG**GCTACATTCTATGTAGCTCTCGTTTAAGAGCTATGCTGGAAACA

sgMYC FWD275 GAAAT**TAATACGACTCACTATAG**AGGATGACGGAGGAGAAAGAGTTTAAGAGCTATGCTGGAAACA

sgMYC FWD276 GAAAT**TAATACGACTCACTATAG**ATTGTGATCAAGATAACCAGTTTAAGAGCTATGCTGGAAACA

sgMYC FWD277 GAAAT**TAATACGACTCACTATAG**AATAGCCTGGTGAATGAGAAGTTTAAGAGCTATGCTGGAAACA

sgMYC FWD278 GAAAT**TAATACGACTCACTATAG**AATTCTAATGGGGAAGCAGGGTTTAAGAGCTATGCTGGAAACA

sgMYC FWD279 GAAAT**TAATACGACTCACTATAG**GCTAGAACAGACCCAATTACAGTTTAAGAGCTATGCTGGAAACA

sgMYC FWD280 GAAAT**TAATACGACTCACTATAG**GTACCTGCCTGGTTGCTTGAGTTTAAGAGCTATGCTGGAAACA

sgMYC FWD281 GAAAT**TAATACGACTCACTATAG**GGATGATGACTCAGAGCGTTTAAGAGCTATGCTGGAAACA

sgMYC FWD282 GAAAT**TAATACGACTCACTATAG**GGAAGATTGCGTCTTCTCCAGTTTAAGAGCTATGCTGGAAACA

| | |
|---|---|
| sgMYC FWD283 | GAAAT**TAATACGACTCACTATAG**AATCAGTTAGGATGCAACCAGTTTAAGAGCTATGCTGGAAACA |
| sgMYC FWD284 | GAAAT**TAATACGACTCACTATAG**GCATCTCAGAAGACCTGGTTTAAGAGCTATGCTGGAAACA |
| sgMYC FWD285 | GAAAT**TAATACGACTCACTATAG**GTCTCCCCCTCTCCCATAGCAGTTTAAGAGCTATGCTGGAAACA |
| sgMYC FWD286 | GAAAT**TAATACGACTCACTATAG**GCCCACCCATGGGTCTCCCCCAGTTTAAGAGCTATGCTGGAAACA |
| sgMYC FWD287 | GAAAT**TAATACGACTCACTATAG**GCAGTCCAAACTGTTTCACCAGTTTAAGAGCTATGCTGGAAACA |
| sgMYC FWD288 | GAAAT**TAATACGACTCACTATAG**GGTCACATTGATCCCACCAGTTTAAGAGCTATGCTGGAAACA |

**Bold**, T7 promoter; underlined, sequence of sgRNA target; FWD, forward primer.
